# JEV helicase targets host MTOC and participates in the organization of pericentriolar viroplasm

**DOI:** 10.64898/2026.07.23.740276

**Authors:** Himanshi Amita, Pranay Eknath Dudhe, Manish Yadav, Brohmomoy Basu, Sudhanshu Vrati, Karthigeyan Dhanasekaran

## Abstract

Flaviviruses contribute significantly to the global disease burden and are known to remodel host organelles to their advantage. Centrosomal microtubule-organizing centres (MTOCs) having established role in cell division and signalling are also targeted by viruses. However, it remains uncertain whether they act as bystanders or actively engage in viral processes. Here, we elucidate the centrosome and cytoskeletal involvement during Japanese Encephalitis Virus (JEV) infection. Our virus-free expression studies revealed a centrosome-targeting region within the C-terminal helicase domain that mediates the association of JEV-NS3 with host MTOCs. When expressed exogenously, JEV-NS3 formed pericentriolar aggresomes resembling the helicase-containing viroplasm distribution pattern of infected cells. These pericentriolar viroplasm appears in the non-neuronal cell types with centriolar consequences while the neuronal cell types fail to show such phenotypes. Centriole depletion assays revealed the proviral role of centrosome, where the viroplasm organization depends on centrosomal MTOCs, and vimentin cages. Additionally, microtubule disruption and dynarrestin blockade assays emphasized the roles of microtubules and dynein in concentrating NS3-containing vesicular packets towards centrosomes, highlighting the centrosome-cytoskeleton axis as a potential target for intervention.

**Summary:** This study demonstrates that JEV helicase is directed to centrosomes via its CTHD domain, using them to initiate pericentriolar viroplasm formation. It also highlights the unappreciated function of centrosomes as a proviral hub, orchestrating cytoskeletal remodelling and viral factory organization depending on the MTOC status of the cell type. These findings highlight the potential host factors that can be targeted for broad antiviral outcomes across flaviviruses.

**Graphical abstract:** 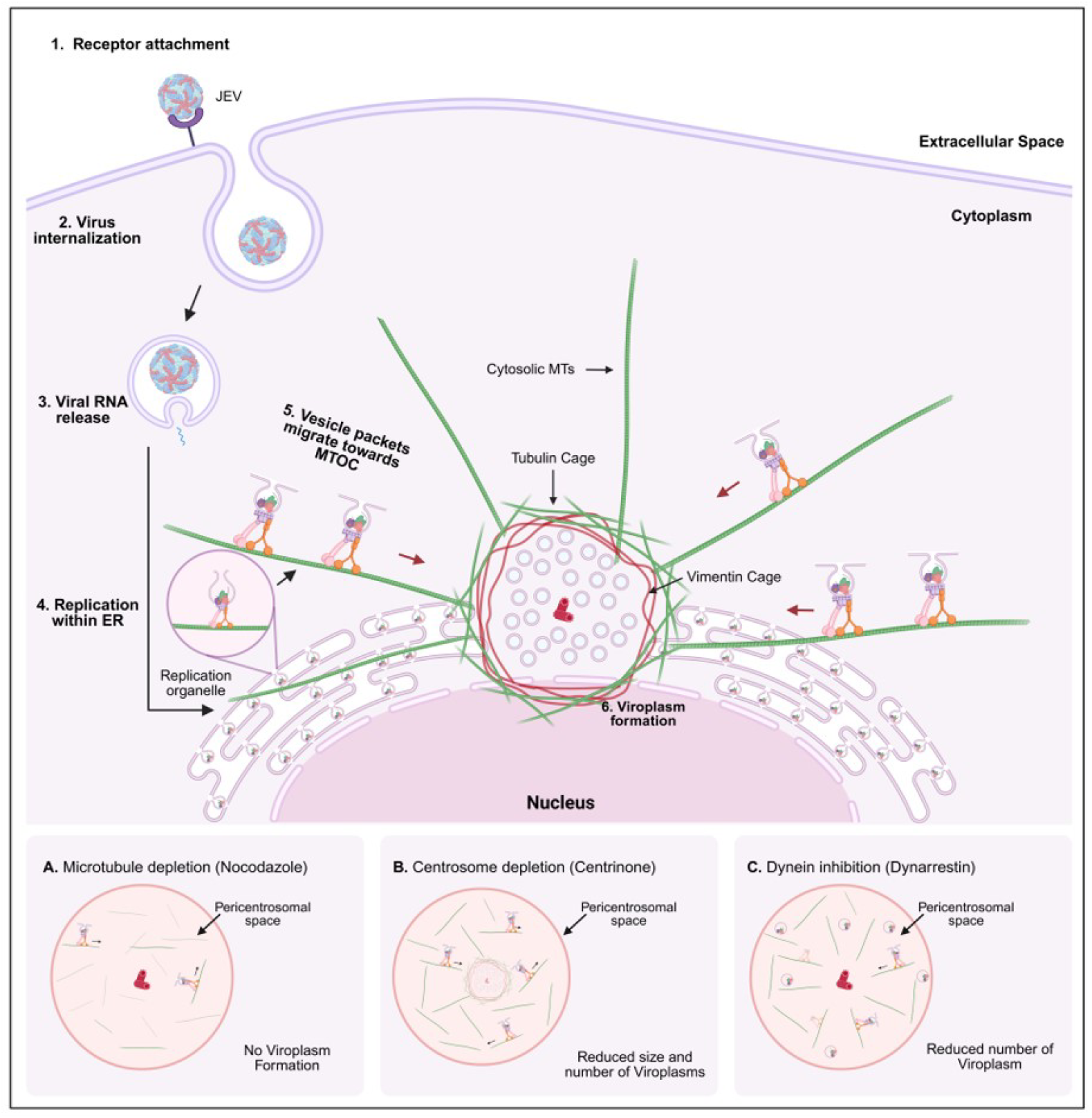

*JEV helicase is targeted to centrosomes to form viroplasm with the aid of microtubule and motor proteins.* JEV lifecycle from receptor binding **(Step 1)** to endocytic internalization **(Step 2)** followed by uncoating and release of viral genome **(Step 3)** followed by replication within the ER derived Vesicular packets **(Step 4)**. This vesicle packets harbors viral proteins that form replication complex, like helicase and replicase and the replicative intermediate, dsRNA. These vesicle packets with the aid of retrograde motor protein gets targeted towards the centrosome **(Step 5)**. Ultimately multiple vesicle packets accumulate in the pericentrosomal region, where it forms a separate compartment enclosed by vimentin and tubulin cage (**Step 6)**. Inset A, B and C depict the effect of cytoskeletal perturbations on Viroplasm formation. **A)** Disruption of microtubules using Nocodazole completely abolishes pericentriolar viroplasm organization. **B)** Centrinone mediated centrosome depletion markedly reduces both the number and size of pericentriolar viroplasm. **C)** Dynarrestin mediated retrograde microtubule transport significantly decreases the reorganization of vesicle packets to form the viroplasm in the pericentriolar region.

## Introduction

Viruses have always relied on the host’s organellar and molecular machinery for their survival, replication, and propagation. While several viral proteins have proven valuable for drug design, host-directed antivirals offer certain advantages over traditional antivirals and are equally important strategies to thwart infection (He et al., 2024). Although numerous instances have been documented to highlight the involvement of the microtubule-based cytoskeleton and its nucleator, organizer, and anchor, termed the centrosome (Amita et. al., 2025; Zhang et al., 2015; Lehmann-Che et al., 2007; Sfakianos et al., 2003), little is understood in terms of Japanese Encephalitis virus infected cells.

Flavivirus related findings have revealed substantial evidence of the virus targeting its host centriolar and cytoskeletal elements. Of these, the Zika virus (ZIKV) replicase (NS5) and helicase (NS3) target the centrosome and cilia, and the consequences of such host–virus interactions in terms of foetal anomalies are well established (Souza et al., 2016; Ticconi et al., 2016; Kesari et al., 2020; Kodani et al., 2022). No equivalent reports exist for Dengue virus (DENV) helicase, although centrosome disruptions occur indirectly via virus-induced microtubule changes or proteostatic stress (Peña J and Harris E, 2011; Chen et al., 2008). In one of the studies by Khadka et al., the association of DENV replicase NS5 in the mitotic stages indicated a phenotype resembling mitotic spindle association rather than a discreet centrosomal association as claimed in the study (Khadka et al., 2011). Moreover, these interactions have not been validated or commented on their association in the interphase stages of the cell cycle. Although it is established that the DENV helicase may exert an indirect influence on centrosomes through mechanisms such as microtubule modulation, the formation of replication organelles on ER membranes near MTOCs (Welsch et al., 2009), or via the cytokine response of flavivirus-infected hosts leading to the expression of interferon-alpha (IFN-α), which results in spindle mispositioning and supernumerary centrosomes as a general response, there has been no direct binding or association of the helicase reported till today (McDougall et al., 2019; Wolf et al., 2017). Similarly, in the case of the JEV-NS3 it localizes primarily to the perinuclear ER membranes (Edward and Takegami, 1993), and to some extent within the nucleus (Uchil et al., 2006) and beyond that it is known to interact with TSG101 and microtubules, but none of these observations suggest a direct centrosomal association (Chiou et al., 2003). In all these instances, it remains unclear whether the centrosome is merely a spectator or if they can actively regulate and dictate the viral subcellular dynamics.

In this study, we attempted to determine whether DENV and JEV helicases can target the centrosome independently, like ZIKV helicase, and further characterized the targeting towards the centrosome and its importance, specifically in the context of JEV propagation. We identified that centrosome targeting is achieved by means of the C-terminal Helicase domain (CTHD) of JEV-NS3 and might be true for other similar flaviviruses that utilize the host MTOC, such as ZIKV. Furthermore, we documented the microtubule-based cytoskeletal aid in organizing the JEV viroplasm, similar to the ZIKV viroplasm, which is known to form toroidal or spherical viroplasms preferentially in the centrosomal vicinity (Buchwalter et al., 2021). The pericentriolar JEV viroplasm reported in this study is heavily dependent on the centrosome and its MTOC function. These viroplasms are restricted to the pericentriolar space with the help of microtubule reorganization and vimentin cage formation to support the organization of JEV viral factories in infected HeLa cells. Our investigations revealed the clustering of several individual vesicular packets derived from the ER membrane, which ultimately formed JEV viroplasms in the pericentriolar vicinity. In contrast, in the case of huh-7 cells the viroplasm showed toroidal organization around the centrosome, and in the neuronal cell type, such as SH-SY5Y, the individual vesicle packets were documented to have preferential MTOC polarization. Furthermore, this study documents the impact of MTOC targeting on centrosome structural and functional integrity and uncovers the importance of centrosomes and their associated cytoskeletal machinery in regulating the aggresome dynamics of viral elements. Our research provides mechanistic insights into the MTOC targeting of viral factories and establishes centrosomes as proviral organelles that promotes JEV infection.

## Results

### JEV NS3 localizes to centrosomes with the aid of its C-terminal helicase domain

To test the centrosomal localization status of flaviviral helicases, we created SNAP tag fusion product-expressing clones for DENV-NS3 and JEV-NS3 proteins, in addition to ZIKV-NS3, which is known to localize to centrosomes, and the related helicase from a non-flaviviral member, HCV-NS3, as a negative control. Individual SNAP tag proteins were labelled with fluorophore tag substrates and combined with immunofluorescence for centrosome labelling. Their subcellular localization was assessed using confocal microscopy upon exogenous expression in HEK293T cells (Figure 1A). Both DENV and JEV helicases were observed to be associated with the centrosome based on the percentage colocalization score, which was comparable to that of ZIKV helicase (Figure 1B), indicating a centrosomal pool of these proteins. In contrast, HCV-NS3 labelling was comparable to that of the SNAP tag alone, which failed to show any such centrosome-localizing preference. Furthermore, the centrosomal localization of SNAP-JEV NS3 was confirmed in HeLa-M cells (Figure S1A) and the expression status at the protein level for all the SNAP tag proteins were established by means of western blot analysis (Figure S1B-E). To identify the minimal and essential fragment responsible for the centrosome targeting of JEV-NS3, additional truncation mutants either lacking the C-terminal tail (CTT), or lacking both CTT and CTHD, or lacking CTT, CTHD, and DEAD domains, were created as N-terminal SNAP-tag fusion products (Figure 1C). These truncated proteins and full-length JEV-NS3 were exogenously expressed in HeLa-M cells to assess their ability to localize to the centrosome (Figure 1D). Comparing the co-localization status of these truncated versions with full-length JEV NS3 revealed that the truncation products lacking the CTHD domain showed a drastic reduction in the centrosome-localizing pool (Figure 1E). Taken together, these results suggest that CTHD is essential for JEV NS3 centrosome targeting. To further assess whether the CTHD domain is also the minimal fragment for centrosome targeting, CTHD alone and the CTHD+DEAD domain were expressed in HEK293T cells, as they were more toxic when expressed in HeLa-M cells (Figure 1F). Their centrosome localization status compared to full-length JEV-NS3 indicates that the CTHD+DEAD domain retains the centrosome localization ability similar to the full-length NS3, while the CTHD alone is capable of co-localizing with the centrosome but slightly compromised as compared to the full length, which indicates that the CTHD fragment of the helicase is both the minimal and essential region for centrosome targeting of JEV-NS3 (Figure 1G). To corroborate this observation, we created a chimeric version of a non-centrosome-localizing HCV-NS3, in which the CTHD segment alone was replaced with JEV-CTHD, to assess its localization status in HEK293T cells (Figure 1H). As expected, we observed that wild-type HCV-NS3 did not localize, whereas the chimeric helicase showed centrosomal localization comparable to that of JEV-NS3 (Figure 1I and 1J). Collectively, these findings indicate that the centrosomal targeting of JEV helicase is primarily facilitated by molecular signatures predominantly located in the CTHD region of JEV NS3 (Figure 1K).

**Figure 1.**
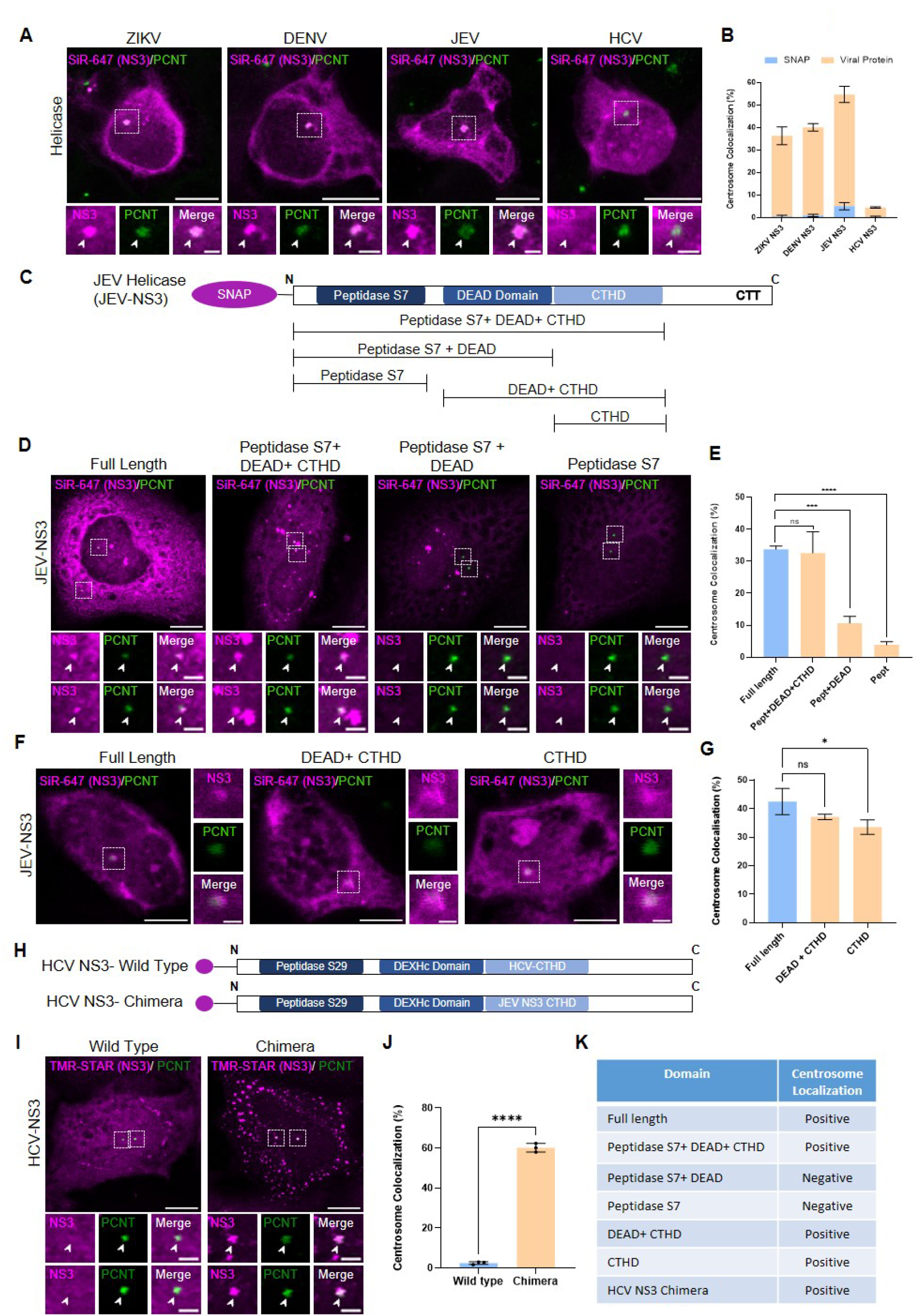
JEV helicases localize to centrosomes with the help of CTHD domain. (A) Confocal micrographs showing the centrosome localization status of indicated helicases in HEK293T cells. (B) Quantification of the number of cells exhibiting centrosome localizing helicases in HEK293T cells (ZIKV NS3, p value- 0.0009, ***; DENV NS3, p value-<0.0001, ****; JEV NS3, p value- 0.0003, *** and HCV NS3, p value- 0.1000, ns), in comparison to empty vector expressing SNAP only. (C) Schematic showing domain organization of full-length JEV-NS3 and the truncations created. (D) Confocal micrographs of the indicated C-terminal domain truncated constructs showing their centrosome localization in HeLa-M cells and (E) quantified for the centrosome localization in comparison to full length JEV helicase (Pept + DEAD + CTHD, p value- 0.9561, ns; Pept + DEAD, p value- 0.0001, ***; Pept, p value- <0.0001, ****). (F) Confocal micrographs of N-terminal domain truncated constructs showing their centrosome localization ability in HEK293T cells and (G) quantified for the number of cells exhibiting centrosome localization in comparison to full length (DEAD + CTHD, p value- 0.1269, ns; CTHD, p value- 0.0209, *). (H) Schematic showing domain organization of the Wild type and Chimeric HCV helicases. (I) Immunofluorescence staining showing their centrosome association of the indicated HCV helicase with (J) quantification for the number of cells exhibiting centrosome localization (HCV NS3 Chimera, p value-<0.0001, ****). (K) Table summarizing the indicated domain constructs and their centrosome localization status. All the datasets were acquired from biological triplicates considering ≥100 individual cells in each set; N=3. The centrosomes are indicated by white arrowheads and marked by Pericentrin in green while the helicases are marked in magenta (Scale bar- 10 μm). Inset represents zoomed in view (Scale bar- 2 μm). All the quantifications represent the mean values with error bars representing the SD values.

### JEV-helicase NS3 clusters in the pericentriolar region with the aid of microtubules

To corroborate the findings from the SNAP tag, a C-terminal HA tag fusion construct was evaluated in HeLa-M cells for their localization status using an immunofluorescence assay (Figure 2A), where the centrosomal localization of JEV NS3-HA was comparable to that of the SNAP tag version of JEV NS3. In addition, the punctate expression pattern of HA tag NS3 it also exhibited a vesicular staining pattern analogous to that of JEV infected membranes derived from the ER that is appreciated in the Figure S2A. In light of the observations derived from a virus-free protein expression system, it is essential to examine the subcellular context of centrosomal association and the role of microtubules in the context of JEV infection. Hence, a comparative analysis focusing on the localization pattern of NS3 in mock infected versus JEV-infected (MOI-5) HeLa-M cells was performed (Figure 2B). These results showed significant pericentriolar accumulation of NS3 in approximately 50% of the infected population, comparable to the association documented in our virus-free protein expression system (Figure 2C). However, none of the infected HeLa-M cells in our experiments showed the direct microtubule-associated pattern of NS3 that was previously reported in BHK-21 cells with infection as well as exogenous NS3 expression (Chiou et al., 2003) (Figure S2B). Moreover, when we attempted to block the proteasomal degradation machinery using MG132 in NS3-HA expressing HeLa-M cells to enrich the centrosomal pool and subsequently followed the localization status by immunofluorescence assays (Figure 2D), we observed a predominant clustering of NS3 preferentially in the pericentriolar region of MG132 treated population when compared to the DMSO (vehicle only) treated cells. This localization pattern was similar to the aggresomes (Johnston et al, 1998; McNaught et al., 2002) that are expected to localize in the pericentriolar space (Figure 2E). Moreover, the radial distribution quantifying the pericentriolar distribution of NS3-HA in the DMSO and MG132 treated population revealed an event of redistribution of the cytoplasmic pool towards the MTOC, (Figure 2F) while the protein expression status of DMSO vs MG132 treated population showed a slight drop (Figure S2C-D) as opposed to the increased stability expected following a proteasomal block that is likely due to the combined toxicity of viral protein and MG132 treatment. Put together, the NS3-HA localization following MG132 treatment reflects a dynamic reorientation facilitated by the centrosome leading to its accumulation in the pericentriolar space like aggresomes which is likely to dependent on the microtubules emanating from the centrosomal MTOCs.

**Figure 2.**
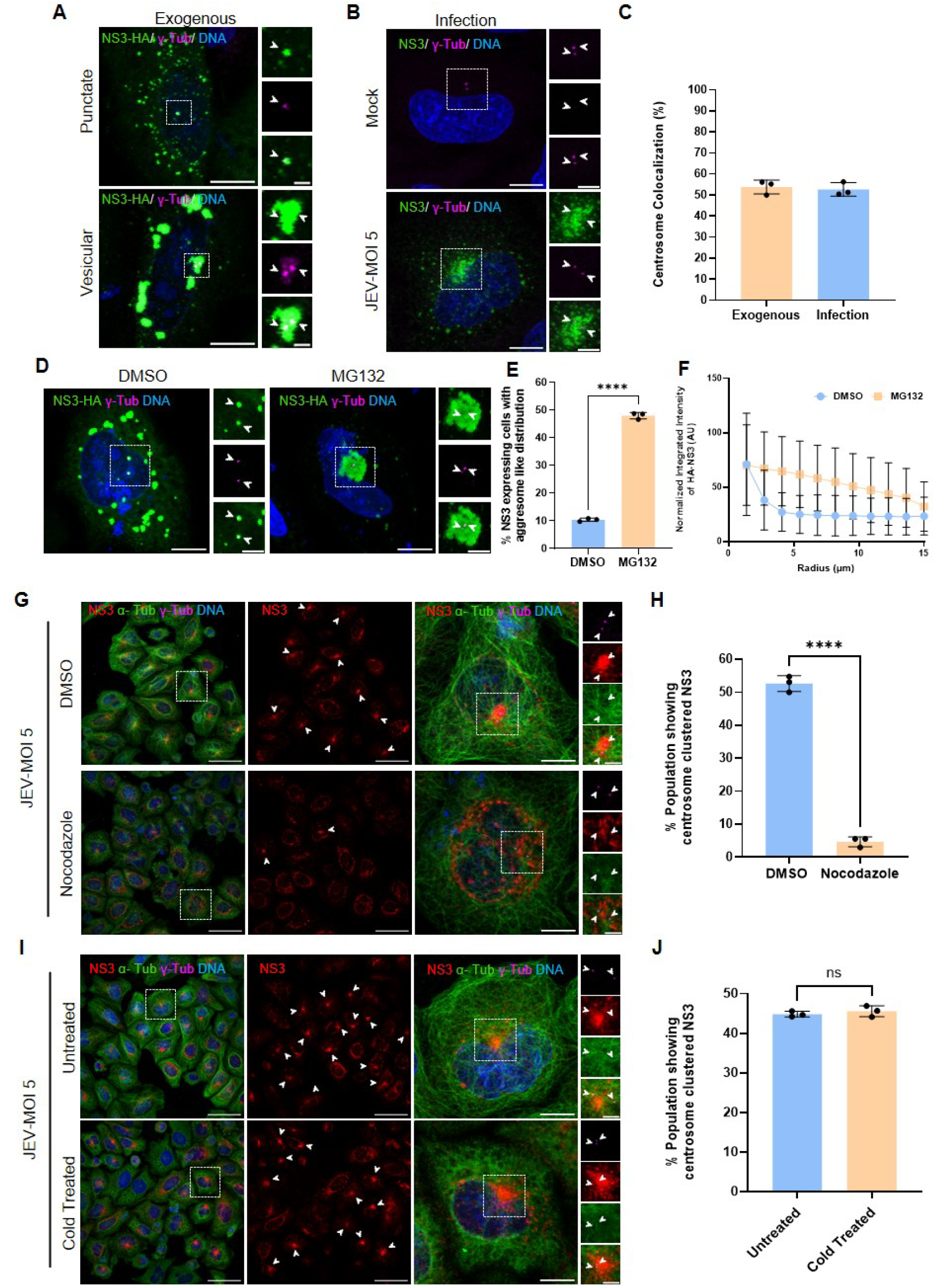
JEV Helicase preferential clustering in the pericentrosomal space depends on microtubules. (A) HeLa-M cells expressing NS3-HA colocalized with the centrosomes in cells exhibiting both punctate and vesicular distribution pattern. (B) Colocalization of NS3 with centrosomes in mock vs JEV infected HeLa-M cells. NS3 is represented in green and γ-tubulin in magenta are represented by white arrowheads while DNA is stained with Hoechst (Scale bar- 10 μm). Inset represents zoomed in view of the colocalized puncta (Scale bar- 2 μm). (C) Quantification comparing the percentage centrosome localizing population in panel A and B (N=3, n≥100). (D) Distribution pattern of NS3 (green) in DMSO vs proteasomal block for 12 hrs with 10 μM MG132 in HeLa-M cells expressing NS3-HA. Centrosome marked by γ-tubulin in magenta are indicated by white arrowheads (Scale bar- 10μm with Zoom in Inset Scale bar- 5 μm). (E) Bar graph of percent cells exhibiting aggresome-like distribution in the indicated population (p value- <0.0001, ****, N=3, n≥100). (F) Radial distribution profile of JEV NS3 comparing between DMSO and MG132 treated cells (N=3, n=25). (G) Confocal micrographs showing NS3 clustering propensity around the centrosomes in the indicated JEV infected population with DMSO and Nocodazole. (H) Quantification of number of cells with pericentrosomal clustered NS3 in DMSO and Nocodazole treated cells (p value- <0.0001,****, N=3, n≥100). (I) Confocal micrographs comparing with and without cold treatment in JEV infected HeLa-M cells. NS3 is represented in red, α-tubulin in green and γ-tubulin in magenta marking the centrosome indicated by white arrowheads (Field view Scale bar- 50μm, single cell view-10 μm and zoom in Inset, scale bar- 5 μm) in panel G and I. (J) Bar graph quantifying the percent of cells with pericentrosomal clustered NS3 in the indicated population (p value- 0.4654, ns, in comparison to untreated cells, N=3, n≥100). All the quantification shows the mean values obtained from biological triplicates with error bars representing the SD values. DNA stained with Hoechst appears in Cyan in all the panels.

To evaluate this hypothesis, we examined JEV-infected HeLa-M cells 24 hpi, in the presence of nocodazole for the final 12 hrs of infection to depolymerize the host microtubules. The necessity of an intact microtubule network for the clustering towards centrosomes was assessed by comparing DMSO vs nocodazole-treated populations using an immunofluorescence assay (Figure 2G). As expected, the pericentriolar clustering phenotype was almost completely abolished in the cells exposed to nocodazole (Figure 2H), whereas the DMSO control showed ∼50% NS3 clustering around the centrosomes. In contrast, when microtubule depolymerization was attempted by means of a brief cold shock treatment during the last 30 min of infection, it did not result in any visible alteration in the presence or absence of such cold shock (Figure 2I), where both the untreated and cold-treated populations showed ∼ 45% infected cells with the pericentriolar clustering phenotype. Although the microtubule organization visualized by anti-tubulin antibodies showed a loss of organization in all cold-shocked cells compared to the untreated population, the number of centrosome-clustered NS3 staining patterns did not change (Figure 2J). Such resistance to cold shock towards the 24 hpi endpoint indicates that the viral elements once clustered around the MTOCs, they are retained in the pericentriolar space with some added aid that is resistant to cold shock. This suggests the possibility of an extra support for the clustering components comparable to the aggresomes retained preferentially in the pericentriolar space due to the vimentin cages derived from the host cytoskeletal rearrangement events (Wileman, 2007).

### Host centrosomal MTOC serves as potent JEV viroplasm organizers

To examine whether the clustering of viral elements in the pericentriolar space is indeed limited by vimentin, we stained mock and JEV infected HeLa-M cells with NS3, vimentin, and γ-tubulin to visualize the organization pattern around the MTOCs. As expected, almost all pericentriolar clustered NS3 staining entities were caged by vimentin (Figure 3A). Furthermore, anti-α-tubulin antibody-based staining was performed to visualize the microtubule reorganization of the mock and infected HeLa-M cells, where the pericentriolar clusters limited by vimentin cages were observed to be well accommodated within the microtubule cages (Figure 3B). These observations indicate the possibility that these NS3 pericentriolar clusters are a part of the viroplasmic organization comparable to that of the toroidal and spherical ZIKV pericentriolar viroplasm reported earlier (Buchwalter et al., 2021). In addition, JEV replicates elaborately within the ER-derived vesicular packets (Hase et al., 1990a; Hase et al., 1990b) like most of the flaviviruses (Romero-Brey and Bartenschlager, 2014), and during the later stages of replication, the majority of the viral non-structural proteins, except for NS3 and NS5, are degraded by the ERAD pathway. Furthermore, when viral components are produced beyond the handling capacity of the ERAD machinery, NS proteins accumulate within the dedicated convoluted membrane structures derived from the infected ER membrane, supporting host cell survival for efficient viral packaging and propagation (Tabata et al., 2021).

**Figure 3.**
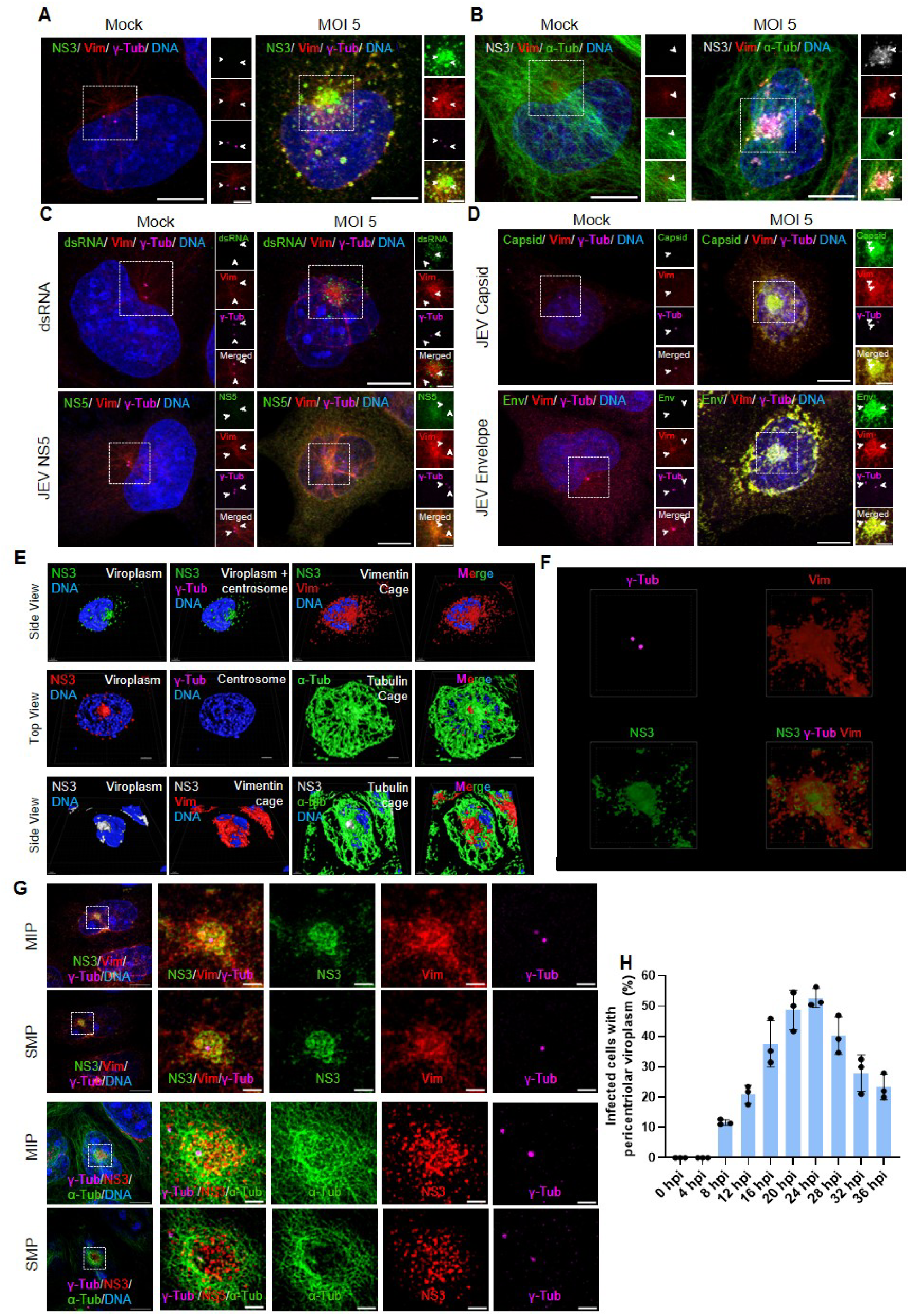
JEV vesicular packets are organized into viroplasm by centrosomal MTOCs in HeLa cells. (A) Vimentin cage encloses centrosome clustered NS3. NS3 is represented in green, vimentin in red and γ-tubulin in magenta are represented by white arrowheads. (B) Confocal micrographs showing clustered NS3 enveloped by both Vimentin and tubulin cages in the infected compared with mock. NS3 is represented in grey indicated by white arrowheads, vimentin in red and α-tubulin in green. (C) Confocal micrographs depicting the co-staining of viroplasm caged by vimentin (red) with components of replication complex like dsRNA or NS5 represented in green, along with centrosomal marker γ-tubulin in magenta indicated by white arrowheads. (D) Confocal micrographs depicting the co-staining of viroplasm caged by vimentin (red) with assembly components either Capsid or Envelope in green, and γ-tubulin in magenta indicated by white arrowheads (Panel A to D, Scale bar- 10μm, Inset represents zoomed in view, Scale bar- 5 μm). (E) 3D rendered confocal micrographs of side and top view of JEV viroplasm marked by NS3 in the context of Vimentin cage and the microtubule rearrangement indicated by α-tubulin in the pericentriolar space with centrosome marked by γ-tubulin antibodies in the indicated color. (F) Snapshot of 3D volume movie generated from the 3D-SIM images of JEV viroplasm in HeLa-M cells with indicated antibodies. (G) 3D-SIM micrographs of JEV viroplasm showing individual vesicle packets marked by NS3 (green) that is enclosed by vimentin cage (red) represented as maximum intensity projection (MIP) or single mid plane (SMP) images in the top panel and the similar image showing the microtubule rearrangement marked with anti α-tubulin antibodies (green) along with NS3 (red) and γ-tubulin (magenta) staining in the bottom panel (Scale bar 10 μm). Zoomed in insets represent the viroplasm (Scale bar 2 μm). (H) Time course experiment plotting the percentage of HeLa population infected with JEV (MOI-5) showing pericentriolar viroplasm (VLS) across the indicated hours post infection (hpi). Data is collected from three independent experiments (N=3, n≥100 infected cells) and plotting the mean values with error bars indicating SD values. DNA stained with Hoechst appears in Cyan in all the panels.

Drawing upon the existing literature on flavivirus, it can be hypothesized that the NS3 clusters organized by the host MTOCs likely function as vesicular packets (VP) that facilitate both the replication and assembly of the JEV. To confirm this replication, components such as dsRNA and NS5 were immunolabelled, revealing a pericentriolar staining pattern similar to that of NS3 (Figure 3C). Similarly, viral assembly components, such as JEV-Capsid and JEV-Envelope, were immunolabelled to reveal a pericentriolar clustering pattern comparable to that of JEV NS3 (Figure 3D). These findings corroborate that the pericentriolar structures observed through NS3 were indeed components of JEV vesicular packets facilitating the viral life cycle within the cytoplasm of the infected host. By integrating the reorganization of vesicular packets by MTOCs within the vimentin cages, we can unequivocally identify these viral component-harbouring pericentriolar structures as distinct JEV viroplasm. These viroplasms supported by vimentin and microtubules were better visualized in 3D upon volumetric rendering of confocal micrographs created from individual stacks captured after labelling the various components of interest (Figure 3E). To further investigate whether the viroplasm is composed of multiple vesicular packets, we conducted immunofluorescence staining to visualize the viroplasm structure using 3D-SIM microscopy, to discern if individual vesicles appear within these viroplasms neighbouring the host centrosomal MTOCs. As predicted, the individual vesicles could be resolved with anti-NS3 staining along with γ-tubulin and vimentin labelling to appreciate the pericentriolar pile-up of these vesicles that are limited by the vimentin cages in the centrosomal vicinity (Figure 3F, 3G and Figure S3A and S3B). To further elucidate the kinetics of JEV viroplasm formation, we conducted a time-course assay to monitor the pericentriolar viroplasm formation in HeLa-M cells infected for various durations (Figure S3C). These outcomes clearly established that the viroplasm was maximally organized around the MTOCs beyond 12 hpi and peaks between 16-24 hpi (Figure 3H). While the involvement of microtubule-organizing centres during infection is acknowledged, it remains uncertain whether the centrosome is directly affected, and if so, to what extent in such a scenario.

### Centriolar structures experience significant alteration during JEV infection

To determine whether there were any structural or functional disturbances in the centrosome itself, we investigated centrosomal integrity in terms of γ-tubulin levels (Figure 4A) to assess structural integrity (Joshi et al., 1992). Our results clearly demonstrate that the centrosomal immunolabelling intensity of γ-tubulin in the infected HeLa-M cells was significantly decreased by ∼25% in comparison to the mock (Figure 4B). In addition, we also examined another centrosomal PCM component Cep152 using anti-Cep152 antibody (Figure 4C) and the centrosomal loading of the same was dropped by ∼32% in the infected population compared with mock population (Figure 4D). Following this, we examined the numerical aberration status of centrosomes upon infection at various time points ranging from 24 to 48 and 72 hpi by tracing the number of γ-tubulin foci that were more than two per cell (Figure 4E). Notably, a significant proportion of the population exhibited supernumerary centrosomes at all time points (Figure 4F), with the highest occurrence documented in the 48 hpi population. Beyond 48 hpi, although there was significant centrosomal amplification, a substantial number of cells were lost owing to the severity of the viral cytopathic effects in HeLa-M cells by the end of 72 hpi. In addition to centrosomes, we assessed whether centriolar satellites (CS), which are considered an important centrosome accessory structure lying in the pericentrosomal space, were affected. We investigated the localization status of CS in mock- and JEV-infected HeLa-M cells using anti-PCM1 antibody-based immunofluorescence (Figure 4G). Strikingly, the PCM1 puncta that appeared in the pericentrosomal region of the mock were almost absent in majority of the infected cells. The same was quantified to determine the overall decrease by ∼65% in PCM1 levels (Figure 4H). Moreover, the preferential pericentriolar distribution pattern observed in the mock population was absent in the infected population (Figure 4I). Since CS dynamics is expected to depend on the microtubules nucleating from centrosomal MTOCs, we examined the infected HeLa-M cells for their microtubule nucleation ability. The same was evaluated by means of the microtubule re-nucleation after a brief cold shock mediated disruption of the microtubule fibres in the infected and uninfected cells using anti-α-tubulin based immunostaining followed by confocal microscopy. Images acquired for the 1 min recovery point from cold shock are likely to allow microtubule re-nucleation, which is revealed by the appearance of spindle aster emanating from the MTOC marked by the centrosomal protein using anti-Cep152 in the mock population, as expected. In contrast, JEV-infected cells showed a significant decrease in the microtubule nucleation ability of centrosomes (Figure 4J). The re-nucleation status was further quantitatively analysed across 1 min post recovery time points which showed nearly 80% drop in the microtubule asters appearing in the infected compared to mock (Figure 4K). Ultimately in the later 4 min recovery time point the microtubule cytoskeleton were re-established in both the mock and infected with a slight drop in the cytoplasmic microtubule filaments appearing in the infected population by ∼14% compared to mock thereby indicating a retardation in the microtubule dynamics rather than a complete blockade of MTOC function during JEV infection (Figure 4L). Eventually, all the analysis pertaining to centriolar phenotypes observed in the infected population clearly demonstrates that JEV infection results in substantial structural and functional disruption of the host centriolar MTOCs.

**Figure 4.**
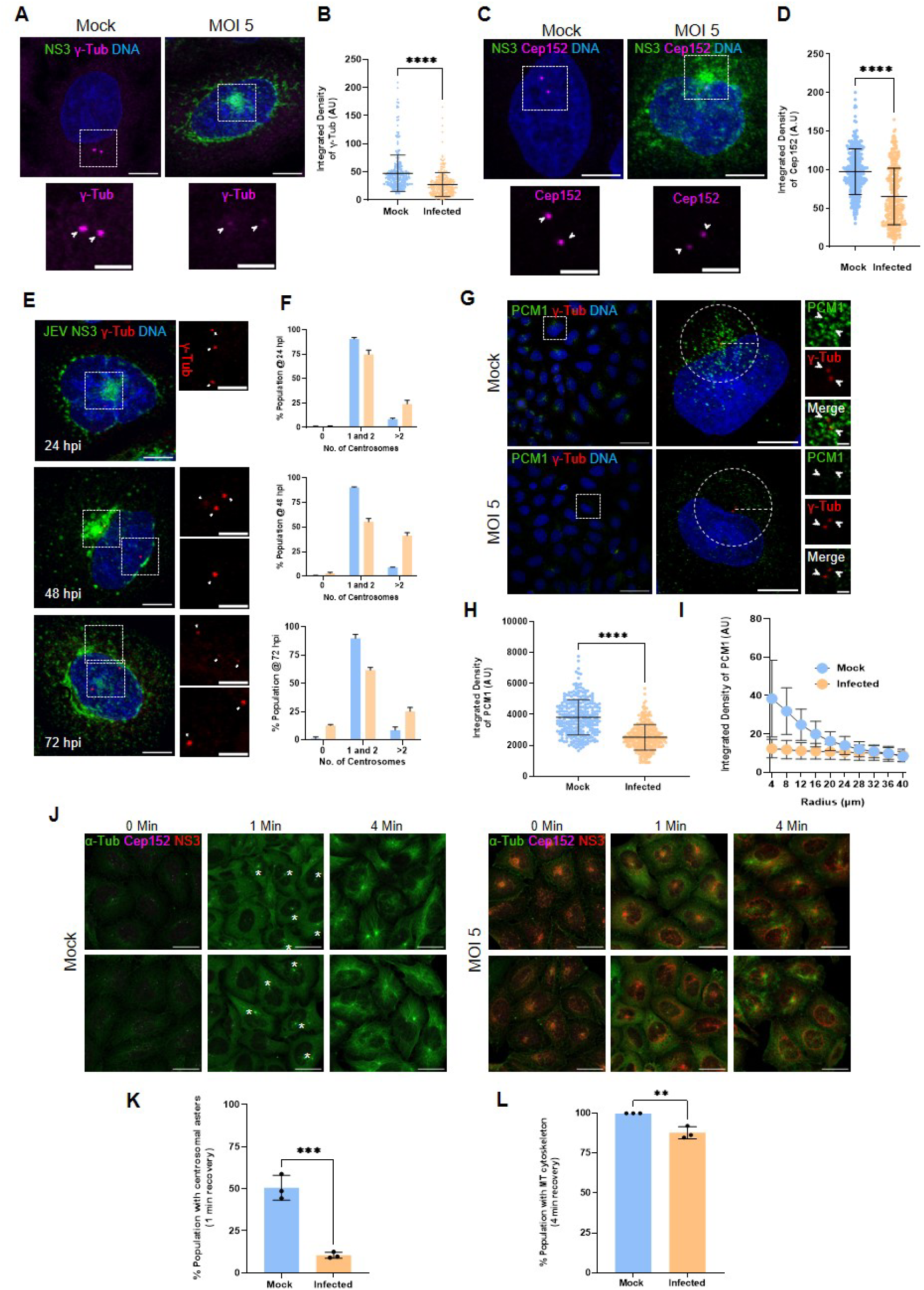
Centriolar implications following JEV infection. (A) Structural aberrations pertaining to JEV infection. Anti-NS3 antibody-based staining (green) is used to identify infected cells and anti-γ-tubulin antibody (magenta) to score the structural aberration in centrosome by scoring the centrosomal fluorescence intensity from the confocal micrographs. Arrowheads indicate centrosomes (Scale bar- 10μm, Inset represents zoomed in view of pericentriolar viroplasm, Scale bar- 5 μm). (B) Quantification for the integrated intensity of γ-tubulin post JEV infection (p value- <0.0001, ****, in comparison to uninfected cells, N=3, n=100). (C) Confocal micrographs of Mock vs infected HeLa-M cells co-stained with anti-NS3 (green) and anti-Cep152 (Magenta). White arrowhead represents centrosome. (D) The centrosomal pool of Cep152 is quantified and represented as a dot plot indicating the mean obtained from three independent experiments where the error bar represents SD (p value-<0.0001,****, N=3, n=100). (E) Numerical aberrations pertaining to JEV infection for 24, 48 and 72 hpi with NS3 (green) and γ-tubulin (red) staining in the confocal micrograph. Arrowheads point centrosomes (Scale bar- 10μm, Inset represents zoomed in view of pericentriolar viroplasm, Scale bar-5μm). (F) Quantification for the percentage of cells with amplified centrosomes post JEV infection at 24, 48 and 72 hpi (N=3, n≥100 infected cells). (G) Distribution of centrosomal satellites post JEV infection in HeLa M cells marked by anti-PCM1 in green and γ-tubulin is marked in red with the white arrowheads pointing the centrosomes (Scale bars for Field view- 50μm and Single cell- 10μm, Zoom in Inset- 2μm). (H) Quantification for the integrated intensity of pericentriolar PCM1 in mock and JEV infected (MOI-5) HeLa-M cells (p value- <0.0001, ****, N=3, n=100). (I) Radial distribution profile of PCM1 in mock versus JEV infected (MOI-5) HeLa-M cells from three independent experiments quantifying 30 cells in each, N=3, n=25. (J) Microtubule re-nucleation assay comparing mock versus infection. Microtubule nucleation was traced at 0 min, 1 min and 4 min after cold shock recovery using anti α-tubulin (green) with anti Cep152 (magenta) marking the MTOC and anti NS3 (red) to identify the infected population. (Scale bar -30 μm). DNA stained with Hoechst appears in Cyan in all the panels. (K) The microtubule re-nucleation enumerated by the number of asters formed (marked by *) in the 1 min recovery time point and plotted as a bar chart (p value- 0.0008, ***, N=3, n≥100 cells). (L) The microtubule cytoskeleton reorganization scored by the reappearance of cytoplasmic microtubule fibers visually in the 4 min recovery time point confocal micrographs and plotted as a bar chart (p value- 0.0054, **, N=3, n≥100 cells). Data represented in panel K and L are the mean values with error bars indicating SD values.

To determine whether these centrosomal phenotypes are universal, we attempted to trace centriolar aberrations in Huh-7 and SH-SY5Y cells, representing non-neuronal and neuronal cell types, respectively. Interestingly, when we infected Huh7 cells, we observed a toroidal organization of JEV viroplasm that assembled around the centrosome (Figure 5A), unlike the spherical organization seen in HeLa-M cells. Upon quantification, approximately 45% (Figure 5B) of the JEV-infected population showed this pattern of organization. In these JEV-infected cells, we noticed that the centrosomal pool of γ-tubulin (Figure 5C) was also decreased by ∼24% compared to the mock (Figure 5D), which is comparable with the observations made in HeLa-M cells. Likewise, we also observed that the supernumerary centrosomes appeared in significant numbers at 24, 48, and 72 hpi within the JEV-infected Huh-7 population compared to the mock-infected population (Figure 5E). In SH-SY5Y cells, no such pericentriolar viroplasm organization was observed; instead, individual vesicle packets were clearly visible as distinct entities and predominantly polarized towards the centrosomal MTOC (Figure 5F). Approximately 60% of these VPs displayed strong MTOC polarization (Figure 5G). Moreover, the centrosomal pool of γ-tubulin (Figure 5H) was also unaltered in the JEV-infected SH-SY5Y cells compared to the mock (Figure 5I), and no signs of centrosomal amplification were documented in the 24, 48, and 72 hpi populations (Figure 5J). The lack of dramatic centriolar phenotypes in SH-SY5Y cells compared to non-neuronal cell types could be attributed to the absence of pericentriolar viroplasmic organization in the neuronal cell type. All together these findings suggest that centriolar MTOCs play an active role in JEV replication and may not be mere bystanders; rather, they may be capable of dictating the course of infection depending on the cell type-specific MTOC status.

**Figure 5.**
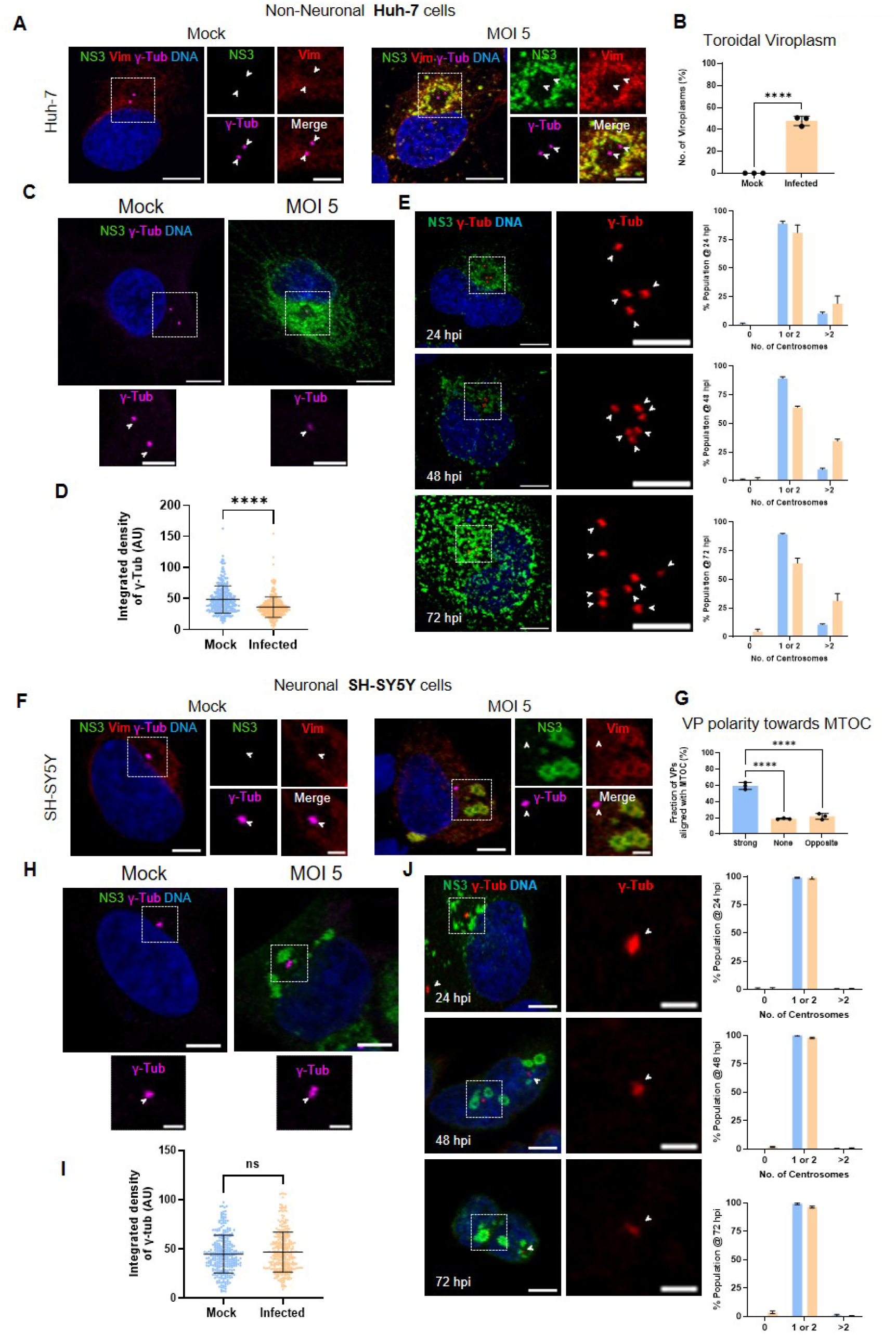
Centriolar consequence comparison between the non-neuronal and the neuronal cell types. (A) Confocal micrographs of mock vs infected Huh-7 cells showing the viroplasm positioned toroidally around the centrosome in the pericentriolar space. (B) Quantification of the toroidal viroplasm appearing in the mock and infected Huh-7 population (p value-<0.0001, ****, in comparison to uninfected cells, N=3, n=100). (C) Structural aberration in JEV infected Huh-7 cells. Anti-NS3 antibody-based staining (green) is used to identify infected cells and anti-γ-tubulin antibody (magenta) to score the structural aberration in centrosome by quantifying the centrosomal fluorescence intensity from the confocal micrographs. Arrowheads indicate centrosomes (Scale bar- 10μm, Inset represents zoomed in view of centrosome, Scale bar- 5 μm). (D) Quantification for the integrated intensity of γ-tubulin post JEV infection (p value- <0.0001, ****, in comparison to uninfected cells, N=3, n=100). (E) Numerical aberrations pertaining to JEV infection for 24, 48 and 72 hpi with NS3 (green) and γ-tubulin (red) staining in the confocal micrograph. Arrowheads point centrosomes (Scale bar- 10μm, Inset represents zoomed in view of pericentriolar viroplasm, Scale bar-5μm). The quantification for the percentage of cells with amplified centrosomes across the corresponding timeline is plotted as percent population with the indicated centrosome numbers (N=3, n≥100 infected cells). (F) Confocal micrographs of mock vs infected SH-SY5Y cells showing multiple VPs polarized towards the MTOC. (G) Quantification of the MTOC polarized VPs appearing in the infected SH-SY5Y population compared to mock (p value- <0.0001, ****, in comparison to uninfected cells, N=3, n≥300). (H) JEV infected SH-SY5Y cells (same as above) to score the centrosomal aberration by quantifying the centrosomal fluorescence intensity of γ-tubulin from the confocal micrographs at 24 hpi. Arrowheads indicate centrosomes (Scale bar- 10μm, Inset represents zoomed in view of centrosome, Scale bar- 5 μm). (I) Quantification for the integrated intensity of γ-tubulin post JEV infection (p value- <0.0001, ****, in comparison to uninfected cells, N=3, n=100). (J) Numerical aberrations pertaining to JEV infection for 24, 48 and 72 hpi in SH-SY5Y cells with NS3 (green) and γ-tubulin (red) staining in the confocal micrograph. Arrowheads point centrosomes (Scale bar- 10μm, Inset represents zoomed in view of pericentriolar viroplasm, Scale bar-5μm). The quantification for the percentage of cells with amplified centrosomes across the corresponding timeline is shown as percent population with the indicated centrosome numbers in the bar chart plotting the mean values with error bars indicating SD values (N=3, n≥100 infected cells).

### Centrosomal MTOCs serve as proviral structures upon JEV infection

Recognizing the involvement of the host MTOC in JEV viroplasm organization, we investigated whether centrosomes function as proviral host components that facilitate the survival, replication, and propagation of JEV upon infection. To evaluate this, we chemically depleted centrosomes using Centrinone in HeLa-M cells (Wong et al., 2015). Subsequently, the centrosome-depleted population was infected with JEV (MOI-5) for an additional 24 h in the presence of Centrinone. This approach was used to assess the status of the JE viroplasm in comparison to the centrosome-retained population, which served as an undepleted control (Figure 6A). The total number of viroplasms decreased drastically by ∼63% in the centrosome-depleted population compared to the control (Figure 6B). Similarly, the distribution status of the JE viroplasm was quantified using the radial distribution profile of NS3 to estimate the radius of the pericentriolar clustered JEV viroplasm, where the centrosome-depleted population showed a drastic reduction in the overall size of the viroplasm by ∼48% compared to undepleted control cells (Figure 6C). Concurrently, we noticed a significant decrease in the overall NS3 staining within the centrosome depleted population; therefore, to authenticate this, a western blot analysis was conducted to compare the expression levels of JEV-Env in both control and centrosome-depleted populations, using the lysates prepared from the respective infected populations (Figure 6D). This analysis demonstrated a ∼ 60% reduction in JEV-Env expression following centrosome depletion (Figure 6E). To rule out the possibility of the direct PLK4 inhibition leading to this viral intervention, HeLa-M cells were subjected to 125 nM Centrinone immediately after infection for the next 24 hrs (Figure S4C) and assessed for the viroplasm formation status (Figure S4D & E) using confocal microscopy and in parallel the lysates were assed for the JEV-Env expression level in the western blot analysis (Figure S4F & G). Both the end points revealed no change in comparison to DMSO control treated population suggesting no direct role for PLK4 in this context. Put together, these findings suggests that the viral events are suppressed in the absence of centrosomes, confirming the proviral role of host centrosomes in JEV infection.

**Figure 6.**
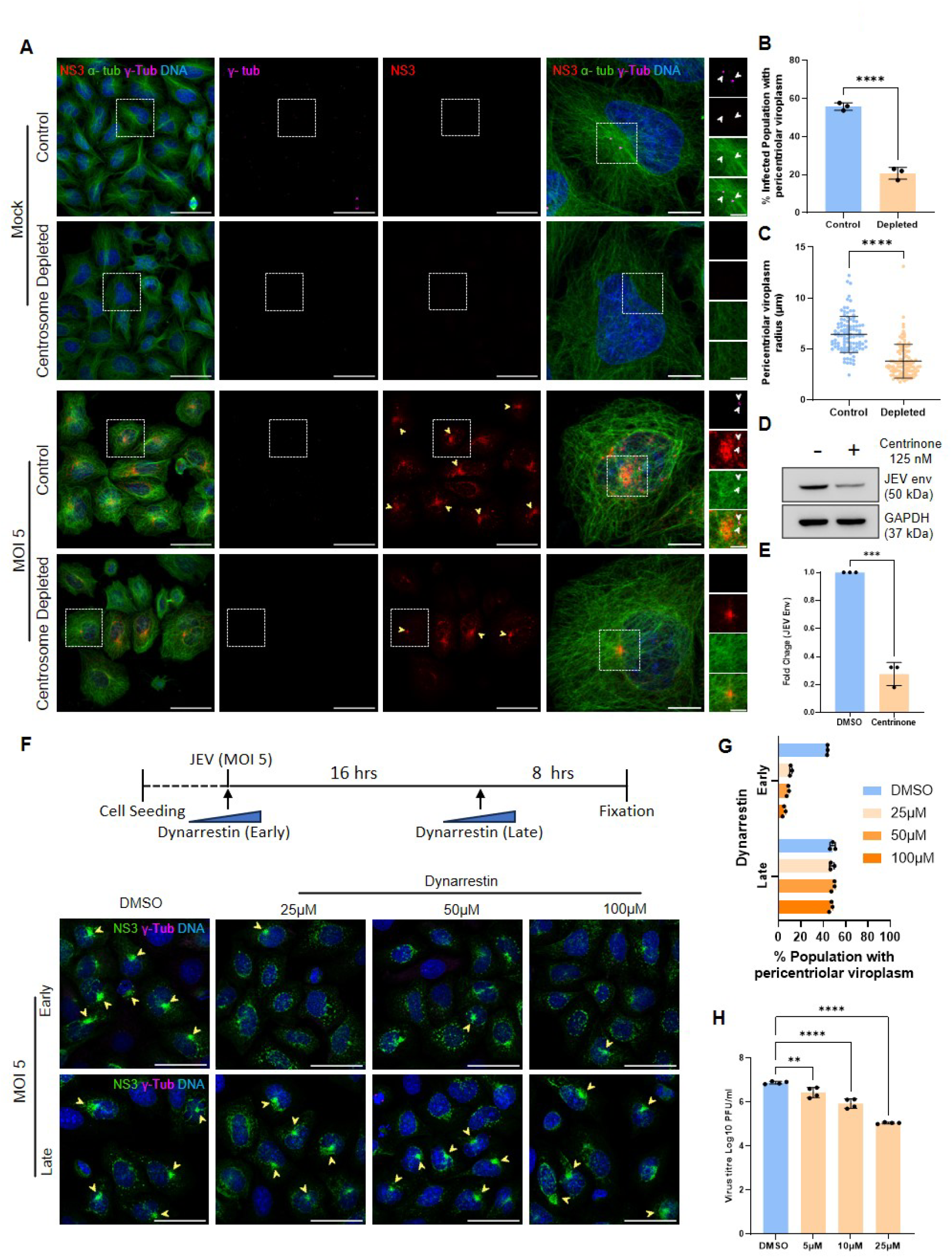
Centrosomes act as proviral organelle supporting JEV propagation. (A) JEV infection 24 hpi confocal micrographs of normal Control (DMSO) and Centrosome depleted (Centrinone) HeLa-M population stained with anti-NS3 (red), anti-α-tubulin (green) and anti-γ-tubulin (magenta) antibodies (Scale bars; Field view -50μm, Single cell -10μm and zoomed in inset showing JEV viroplasm -5μm). (B) Quantification for percentage population having pericentriolar viroplasm in control and centrosome depleted infected cells (p value- ≤0.0001, ****, N=3, n≥100). (C) Quantification of viroplasm radius in the indicated population across three independent replicates measuring 35 cells in each set (p value- <0.0001, ****). (D) Western blot images showing JEV Envelope expression in normal (Centrinone -ve) and centrosome depleted (Centrinone +ve) infected lysates harvested 24hpi. (E) Densitometric quantification of panel D from biological triplicates (p value-0.0001, ***). (F) Schematic of retrograde motor protein block using Dynarrestin treatment and the corresponding confocal micrographs showing NS3 in green and γ-tubulin in magenta after the early and late timepoint treatment of JEV infected HeLa-M cells with the indicated concentrations. Panel A and F, yellow arrowheads indicate the pericentriolar viroplasm while the centrosomes in inset are indicated by white arrowheads. (Scale bars, Field view -50 μm, Single cell -10 μm). Zoom in inset -5 μm). (G) Quantification of percentage cells nucleating pericentriolar viroplasm in Dynarrestin treated compared to DMSO control shown in panel F ((Early: 25 µM p value- <0.0001, ****; 50 µM p value- <0.0001, ****; 100 µM p value- <0.0001, ****, in comparison to untreated cells; Late: 25 µM p value- 0.9542, ns; 50 µM p value- 0.9338, ns; 100 µM p value- 0.6864, ns, in comparison to untreated cells; N=3, n≥100) represented as bar graph plotting the mean values with error bars indicating the SD values. Panel A-G are infected with JEV (MOI-5). (H) Plaque assay to assess the titre of supernatant collected from 0, 5, 10 and 25 μM dynarrestin early timepoint treatment. The viral titer is represented as bar graph of the mean values with error bars indicating SD values (One-way ANOVA Dunnets multiple comparison test with 5 µM p value- 0.0056,**, 10 µM p value- <0.0001, ****; 25 µM p value- <0.0001, ****; N=4). DNA stained with Hoechst appears in Cyan in all the confocal micrographs.

Since our previous observations have established that the host microtubules and their centrosomal organizers (MTOCs) participate actively in some of the flaviviral life cycles, we wanted to check whether blocking this support could suppress the JEV viral events. Hence, we blocked dynein motor-mediated retrograde microtubule transport in HeLa-M cells using dynarrestin (Höing et al., 2017) to assess the viroplasm formation status (Figure 6F-Early). Remarkably, the inhibition of this retrograde transit towards the MTOC significantly diminished the pericentriolar organization status of the JEV viroplasm in a dose-dependent manner while maintaining an intact centrosome in all observed instances (Figure 6G). The majority of viroplasm blockade was accomplished with a minimal dose of 25 μM dynarrestin administered over a 24-hour period. On the contrary, no visible differences were observed in the organization of the viroplasm when challenged with dynarrestin even up to 100 μM when the drug was added beyond the 16 hpi timeline which is already expected to have mature vimentin-caged viroplasms formed in the pericentriolar space (Figure 6F-Late). These observations prompted us to hypothesize that blocking the host motor protein could be an effective host-directed antiviral strategy. Hence, we tested this using increasing doses of dynarrestin following infection and followed the viral titre 24 hpi using the culture supernatants collected from the infected HeLa-M cells in the presence and absence of dynarrestin by means of plaque assays using Vero cells. (Figure 6H). Notably, we observed a drastic dose dependent reduction in the viral titre in comparison to the DMSO-treated population infected with JEV across all the tested dose from 5 to 25 μM dynarrestin proving the proviral role of motor protein that aids in JEV viral propagation. Collectively, our findings underscore the critical role of centrosomes, microtubules, and motor proteins in JEV viroplasm formation and identify potential microtubule-based targets crucial for the intervention of JEV infection. In the future, the insights obtained from JEV and other similar flaviviruses, as well as other RNA viruses that utilize host microtubule-based components, can be collectively harnessed to develop broad-spectrum host-directed antiviral strategies aiming effective combat against such viruses.

## Discussion

Flaviviruses, including Zika, Dengue, and Japanese Encephalitis viruses, exhibit notable similarities in their interactions with host factors during infection (van Leur et al., 2021). The flavivirus life cycle predominantly occurs within the cytoplasmic compartment and extensively involves host organelles, such as ribosomes, the ER, and the Golgi apparatus (Fishburn et al., 2022). Among these three viruses, ZIKV (Buchwalter et al., 2021) and JEV (Xie et al., 2024) have been documented to form viroplasms at certain stages of their life cycle, whereas DENV forms replication organelles purely involving the ER membranes without forming the classical viroplasm (Teo and Chu, 2014). However, all these structures ultimately facilitate efficient replication and assembly to propagate the virus. Although there is currently no consensus regarding the spatiotemporal organization of viral factories across flaviviruses, it is worthwhile to investigate and compare these structures in future studies. Our findings on flavivirus helicases, which can target host centrosomes beyond other cytoplasmic structures, suggest that they may influence the organizational dynamics of their viroplasms. Evidence supporting this hypothesis has been found in JEV-infected HeLa-M cells, where NS3 facilitates the organization of pericentriolar viroplasm during infection. Our findings suggest that in centrosome-dependent flaviviruses, such as JEV, the viral helicase can associate with microtubules and facilitate the trafficking of ER-derived viral factories via dynein motors towards MTOCs. This process ultimately aids in pericentriolar organization, forming a specialized viroplasmic compartment encased by vimentin and microtubule filaments near the centrosomes. Such an organization in proximity to the host MTOC may enhance the efficiency with which the pericentriolar proteostatic machinery is harnessed for viral replication while simultaneously shielding against host antiviral factors through the cytoskeletal shield organized by MTOCs in the pericentriolar space.

Previous research has indicated the potential for DENV and JEV helicases to directly bind to the centrosome, similar to ZIKV helicase. Additional proteomic studies have suggested the involvement of centriolar components in DENV and JEV infections, although these findings have not been validated. For the first time, we conclusively demonstrated that virus-free proteins can associate with the centrosome in the context of JEV and DENV helicases. Furthermore, our domain mapping studies identified the CTHD of JEV helicase as the minimal and essential fragment responsible for targeting flaviviral helicases. Our proteasomal block experiments with NS3-HA expression or the nocodazole block, along with infection traced by anti-NS3 staining patterns, revealed the dependence of JEV on the host microtubule for the pericentriolar clustering of VPs. This clustering event, leading to viroplasm organization, is known to occur in the context of ZIKV infection with a preferential pericentriolar organization (Buchwalter et al., 2021). A similar scenario of viral replication factories has been suggested for DENV (Teo and Chu, 2014), and the same is anticipated for JEV, given the resemblance in the utilization of host machinery among related flaviviruses. A study by Ruibing Cao’s group reported vimentin rearrangement via the CDK1-PLK1 axis, which is mediated by the NS1 and NS1’ proteins, promoting JEV replication (Xie et al., 2024). A follow-up study showed that JEV NS1 aids in concentrating the ER membranes towards the perinuclear region, independent of the host cytoskeleton, ultimately supporting the virus replication organelles (Xie et al., 2025). However, no reports have documented the centriolar context of JEV-infected hosts to date. For the first time, we emphasize that the centriolar MTOC facilitates the organization of JEV viroplasms. Since, it is known that flaviviral helicases are not embedded within the VP membrane, they are able to interact with multiple host components (Lee et al., 2026). Consequently, ER-resident JEV-NS3 is expected to facilitate the trafficking of multiple VPs across microtubules via motor proteins to reach the pericentriolar space, creating a well-organized viroplasm. Moreover, our study indicates that pericentriolar organization benefits the virus. Unlike ZIKV infection, in which viroplasms can form independently of host centrosomes, JEV relies significantly on centriolar MTOCs for viroplasm formation, and the same is affected significantly when centriolar MTOCS are depleted. This conclusion was derived from observations documented upon chemically depleting centrosomes, where the number, distribution, and amount of viroplasm decreased drastically compared to the undepleted population in HeLa-M cells. Further, the JEV viroplasm begins to attain its pericentriolar organization by approximately 12 hpi and peaks by 20 hpi with the aid of host microtubule tracks assisted by dynein motors. Any attempt to disrupt this support beyond the 16 hpi time point showed no effect on the pericentriolar viroplasm, as vimentin caging and microtubule reorganization are already set in motion to maintain the pericentriolar organization. It is possible that NS1 mediated ER remodelling precedes the pericentriolar movement, while CDK1-PLK1 vimentin rearrangement occurs alongside microtubule rearrangement dynamics in the early stages of infection, ultimately supporting pericentriolar viroplasm organization and viral replication.

The viroplasm organization kinetics of JEV documented in this study follow a pattern similar to that of ZIKV, where mature ZIKV viroplasms are observable by 24 hpi (Buchwalter et al., 2021). In contrast, DENV exhibits a delayed pattern, with mature perinuclear replication organelles observed at 48 hpi (Teo and Chu, 2014). Attempts to disrupt microtubules beyond the mature viroplasm formation timeline of JEV (between 16 and 24 hpi) did not affect the pericentriolar organization of JEV viroplasms in HeLa cells. These differences highlight that flaviviruses, such as ZIKV, DENV, and JEV, exhibit a strong propensity to engage the host MTOC; however, they may vary in the timing and extent of their dependence on such centriolar structures. With Japanese encephalitis (Acharya et al., 2026) and other flaviviral infections (Liang and Dai, 2024) emerging as public health threats in certain Asian countries, it is imperative to understand these differences across multiple cell types to consider targeting them for the development of host-directed antivirals in the future.

The structural and functional phenotypes associated with centriolar structures, including PCM disruption, centriolar amplification, centriolar satellite disturbances, and microtubule nucleation blockade in JEV-infected cells, closely resemble the scenario of pericentriolar space-organized aggresomes in response to protein aggregation. These phenomena may either trigger the elimination of the infected host or alter the inflammatory and immune responses of the infected cells to the advantage of the virus, as in the case of HIV-1 (Lacaille and Androlewicz, 2000). A similar loss of PCM1 has also been reported for ZIKV (Wen et al., 2019). Moreover, centriolar amplification has been documented in hosts infected with ZIKV and DENV (Wolf et al., 2017) as well. There remains a chance that these alterations could occur indirectly due to virus-induced stress, or they could be a direct consequence of centriolar interaction facilitating viroplasm organization in the early and mid-stages of the viral life cycle. Whereas during the later stages of the viral life cycle, the centrosomes are deliberately exhausted to prevent the host clearance machinery from reaching the pericentriolar viroplasm. Therefore, it is important to thoroughly examine such possibilities in the future and determine whether these changes have proviral or antiviral consequences.

In addition, comparative studies across other neuronal and non-neuronal cell types indicate that centriolar states vary depending on the cell type, and their involvement in response to viral infection also differs markedly. Huh-7 hepatocarcinoma cells behave similar to HeLa cells, where the viroplasm is organized in the pericentriolar space with a typical toroidal appearance, along with PCM loss and supernumerary centrosomes upon infection. In contrast the SH-SY5Y neuroblastoma cells no such pericentriolar organization was seen. Rather, the viral replication packets appear to be much larger in size and polarized towards the MTOC without building the fused viroplasm that is observed in non-neuronal cell types such as HeLa and Huh-7. This is reflected in terms of the centriolar phenotypic consequences, where PCM loss and centriolar amplification are not observed with a similar timeline of JEV infection in Sh-SY5Y cells. It would be interesting to further study these differences across multiple cell types to understand the holistic contribution of centriolar structures across various tissues involved in JE.

Interference with either centrosomal microtubule nucleation and organization, disruption of the support provided by cytoplasmic microtubule tracks, or obstruction of the motor protein facilitating retrograde transport along the microtubules is sufficient to impede viroplasm formation. In summary, each of these strategies targeting the microtubule or its organizer can effectively interrupt viral replication and propagation. One such demonstration of this concept is the overall reduction in viral titre achieved through dynarrestin treatment. Moreover, the potential for developing strategies targeting these microtubules supported pro-viral events are promising for the creation of broad acting host-directed antivirals against JEV and other related flaviviruses. A report suggesting an antiviral role for tubacin, even when administered prior to infection with JEV, also supports the idea of microtubule acetylation as a putative host-directed antiviral strategy (Lu et al., 2017). Hence, centriolar and cytoskeleton-targeting drugs will be a lucrative option in the future, provided that the host dynamics are minimally altered, for which a deeper insight into the viral spatiotemporal dynamics is crucial to the success of such approaches. Overall, this study uncovers the previously underappreciated role of the centrosome as a dynamic regulatory hub in JEV infection and highlights the centrosome-cytoskeleton axis as a potential target for flavivirus intervention.

### Limitations

Although viroplasmic dynamics are well established in our study, the dynamics of viroplasms within in vivo models, particularly in relevant tissues and cell types during infection, remain inadequately characterized. The anti-JEV-NS3 antibody used in this study is not suitable for performing immuno-pullout studies at present and that limits us from understanding the host cytoskeletal factors directly interacting with them in the course of infection. The HA-tag could be used for this purpose in the future if it is co-incorporated in the viroplasm along with the viral infection across various cell types. Alternatively, a replicon with HA-NS3 needs to be created to pursue such studies in the future. Centriolar phenotypes, such as supernumerary centrosomes, may or may not be a direct consequence of the virus; for instance, it could be a stress-induced or ROS-mediated DNA damage response that can end up in such phenotypes. This must be experimentally deciphered for a better understanding in the future. Similarly, the loss of centrosomal and centriolar satellite components could be a part of the global transcriptional and translational shut-off or could be specific to JEV infection. In addition, the variability of such phenotypes across neuronal and non-neuronal cell types will be an interesting aspect to investigate to understand why certain tissues are more susceptible and permissive to viral pathogenesis in general.

## Material and Methods

### Plasmid, Cloning and Site-directed mutagenesis

ZIKV NS3 (Addgene #79635), DENV-4 NS3 (Addgene #155320) (Phoo et al., 2020), and HCV NS3 (Addgene #17645) (Budhu et al., 2007) were used as templates for subcloning into the pSNAPf-C1 vector (Addgene #58186). All these genes were amplified using Phusion High-Fidelity DNA Polymerase (ThermoFisher). ZIKV and HCV NS3 were cloned into the XhoI and KpnI restriction sites, and DENV-4 NS3 was inserted into the HindIII and XmaI sites. JEV NS3 was commercially synthesized and cloned into the pSNAPf vector (NEB) between the XhoI and NotI sites of the MCS by Synbio Technologies, USA. All C-terminal domain truncations constructs were generated using the Q5 Site-Directed Mutagenesis kit (NEB) following the manufacturer’s recommendation while the N-terminal truncation constructs were subcloned using appropriate primers to amplify the region of interest and subsequently cloned into pSNAPf using XhoI and NotI sites. All the clones described in the study were confirmed by Sanger sequencing. The primers used to generate these clones and mutations are listed in the supplementary Tables S1 and S2 respectively. JEV NS3-HA tag construct is a kind gift from Dr. Manjula Kalia (Sehrawat et. al. 2021).

### Cell culture and Transfection

The following human cell lines were used in this study: HEK293T (ATCC), Huh-7 (Cytion), SH-SY5Y (ATCC) HeLa-M (gift from Dr. Patrick Lusk, Yale School of Medicine, New Haven, CT), C6/36 and Vero cells (a kind gift from Dr. Sudhanshu Vrati, Regional Centre for Biotechnology, Faridabad, India) that were tested to be free from mycoplasma. HeLa-M, Huh-7, SH-SY5Y and HEK293T cells were maintained in DMEM supplemented with 10% Foetal Bovine Serum FBS (HiMedia). Vero cells were grown in MEM (HiMedia) with 10% FBS. All media were supplemented with PenStrep (Gibco). For exogenous viral protein expression studies, cells were seeded in 35 mm culture dishes at a confluency of 0.2 × 10^6^ and transfected using the FuGENE HD transfection reagent (Promega) as per the manufacturer’s protocol.

### SNAP-Cell substrate labelling and Immunofluorescence Assay

A concentration of 0.6 mM SNAP-Cell substrate with 647-SiR/ TMR-STAR conjugate (NEB) was used to label SNAP-tag fusion proteins mentioned in the study. Briefly the cells were incubated with (1:1000 dilution in complete media) SNAP substrate for 30 min at 37°C and 5% CO2. To remove unbound substrate, three washes with complete media were given for 2-3 minutes. This was followed by replacing the complete media for 30 minutes at 37°C and 5% CO2 for destaining and finally fixed at the assay end points after three washes with 1XDPBS to proceed with immunofluorescence assays.

Indirect immunofluorescence assays were performed after fixing the cells with chilled 100% methanol for 5 min at -20°C. Blocking was performed for 1 hr at room temperature (RT) in 5% FBS prepared in PBST (1XDPBS containing 0.1% Triton-X 100) followed by incubation in primary antibody prepared in 1% FBS containing PBST for 1 hr at RT and then washed thrice with 1% FBS containing PBST. Secondary antibody prepared in 1% FBS containing PBST was added to the coverslip and incubated for 1 hr at RT. Unbound antibodies were finally removed by washing with 1X DPBS thrice and stained the DNA with 1:6000 diluted Hoechst 33342 (Sigma). The antibodies and their respective dilutions are mentioned in the supplementary data, Table S3.

### Virus Propagation, Plaque assay and infection

We used the Japanese Encephalitis Virus (strain <u>P20778</u>, GenBank accession no. <u>AF080251</u>) (a kind gift from Dr. Sudhanshu Vrati, Regional Centre for Biotechnology, Faridabad, India) for our study. C6/36 cells were infected with an MOI of 0.01 where the virus was diluted in L-15 media (Himedia) and incubated for 1 hr at 37°C and later replaced with fresh L-15 media containing 10% FBS. The supernatant was collected between 72 and 96 hours post-infection (hpi) and centrifuged at 8000 rpm for 15 min at 4°C and then filtered using a 0.45 μm pore size filter before storing the aliquots at -80°C. Plaque assay was performed to assess the viral titer using Vero cells (a kind gift from Dr. Sudhanshu Vrati, Regional Centre for Biotechnology, Faridabad, India). Briefly 90% confluent Vero cells plated in 12 well culture plates were infected with the serial dilutions of the viral supernatant. The cells were incubated for 1 hour with gentle tilting every 15 minutes. Post infection, the virus was aspirated, and cells were washed with prewarmed 1x DPBS. Later, 1.5 ml of the overlay media (2x MEM and 2% Agarose in 1:1 ratio) was layered on top of the cells to solidify for 30 min and then incubated for 6 days at 37°C. At the end cells were fixed with 1 ml of 3.7% formaldehyde and stained with 0.1% crystal violet prepared with 20% ethanol (1 ml) for 30 minutes, followed by rinsing the plate with tap water and then allowed to airdry overnight to count the plaques and to estimate the viral titer.

For viral infection, the cells were first washed with 1X DPBS. The required viral inoculum (MOI-5) was diluted in DMEM without antibiotic and layered on top of the cells and incubated for 1 hour. After 1 hour, the inoculum was removed, replaced with complete media, and incubated for the indicated time points.

### Drug treatments

For centrosome depletion experiments: To gradually deplete centrosomes, HeLa-M cells were treated with the PLK4 inhibitor, Centrinone (Tocris Bioscience) at a concentration of 125nM for three days, with drug replenishment every 24 hours as opposed to one day treatment for PLK4 inhibition without centrosome depletion. These centrosomes depleted population were harvested counted and seeded according to the requirement either in the presence of centrinone or DMSO as a solvent control. This was followed by JEV infection (MOI-5) and continued to maintain in centrinone or DMSO containing media respectively until the assay endpoint.

Microtubule Depolymerization Assay: Following JEV infection in HeLa-M cells after 12 hpi they were treated with 200nM Nocodazole (Selleck Chemicals) or DMSO for next 12 hrs to achieve depolymerization chemically (Mikhailov and Gundersen, 1998). Cells were then fixed after 24 hrs and proceeded for immunofluorescence assay. For the rapid MT depolymerization assay cold shock was given on ice for 30 min at the end of 24 hpi and fixed immediately in chilled methanol to proceed for immunofluorescence assay.

Proteasome block assay: The proteasome blockade was achieved in HeLa-M cells using 10μM MG132 (Sigma-Aldrich) for the last 12 hrs of the 24 hrs post-transfection. Cells were finally methanol fixed after the end point and proceeded for immunofluorescence assay.

Motor protein block: To block cellular transport machinery, we used a reversible inhibitor of cytoplasmic Dynein 1, Dynarrestin (Sigma-Aldrich) for the indicated dose and time depending on the assay to evaluate either the JEV viroplasm formation or the total virus production by means of viral plaque assays.

### Microtubule re-nucleation assay

Mock and Infected cells were subjected to cold shock briefly over ice for 30 min to depolymerise the microtubules and then released with pre-warmed complete media and incubated at 37°C for the indicated time to allow the re-nucleation event and then subsequently fixed with 4% PFA to visualise the microtubules using anti-α-tubulin antibody based immunofluorescence imaging.

### Western Blot Analysis

For protein expression assessment, the cells were washed with 1X DPBS twice and harvested in RIPA lysis buffer (HiMedia) containing protease (Roche) and phosphatase inhibitor (Roche) at 4°C and further incubated at 4°C with constant agitation for 30 min and then centrifuged at 13000 rpm for 5 min at 4°C. The obtained supernatant was used for western blot analysis, after resolving them using 10% SDS-PAGE and transferring onto 0.2 µm nitrocellulose membrane (BioRad). These blots were incubated for 1 hr in blocking buffer (5% Skim milk powder in TBS and 0.1% Tween-20) followed by primary antibody prepared in TBST (2.5% Skim milk powder in TBS and 0.1% Tween-20) for 1 hr. After washing with TBST thrice, the blots were probed with HRP-conjugated secondary antibody for 1 hr and then visualized with Chemi-luminescent substrate-femtoLUCENT PLUS HRP (G-Biosciences) using Chemidoc. The antibodies and their respective dilutions are mentioned in the supplementary data, Table S3.

### Confocal and 3D-SIM microscopy

Fixed cell confocal microscopy images were acquired either in Leica TCS SP8 laser scanning optical confocal microscope or Leica Stellaris using HCX-PL-APO-CS 63X magnification and 1.4 NA oil-immersion objective and a HyD (hybrid) detector, with 0.2µm step size z-stacks. All image acquisition settings were kept identical for control and test samples. Leica LASX software was used to control various imaging parameters during image acquisition. The images were then processed in ImageJ for scaling and representing the ROIs.

Super-resolution imaging was performed using a Nikon N-SIM S microscope (Nikon Instruments Inc., Japan) equipped with an SR HP Apo TIRF 100×/1.49 NA oil immersion objective lens, a Hamamatsu ORCA-Flash4.0 sCMOS camera, and excitation laser lines at 405, 476, 545, and 637 nm, as appropriate for the respective fluorophores. Images were acquired with a system calibration of 0.065 µm/pixel, and z-stack datasets were collected at a 0.2 µm step size using structured illumination with five phase-shifted excitation patterns at three angular orientations for each optical section. Raw SIM image datasets were reconstructed using NIS-Elements Advanced Research software (Nikon Instruments) with optimized reconstruction parameters to minimize artefacts while preserving super-resolution fidelity. For multicolour imaging, chromatic aberration and channel misalignment were corrected using 80 nm fluorescent bead-based calibration, and the resulting transformation matrix was applied during image reconstruction prior to downstream image analysis.

### Image analysis

ImageJ/Fiji (2.16.0/1.54P; Java 1.8.0_322) was used for image rendering and analysis. Adobe Illustrator was used to arrange figures and to draw the schematics while BioRender.com was used to generate the graphical abstract. For quantification of viral protein colocalization with the centrosome, a line across the centrosome was drawn in software Fiji. RGB profiler plugin was then used to validate the overlapping peaks between the two channels. The cells with overlapping peaks were considered to be positively localizing to centrosome, while rest of the population was considered negative. For estimating the centrosomal intensity, an ROI of 7*7 pixels encompassing the centrosome was used to measure the fluorescence intensity. The same ROI was used to measure the intensity of background, adjacent to the centrosome to obtain the background subtracted fluorescent intensity for the centrosome. These values were exported in an excel file, followed by statistical analysis. To quantify the pericentriolar distribution, a circular ROI was drawn centred around the centrosomal marker. The Radial Profile plugin in Fiji was used to obtain the normalized integrated densities across the radius and exported to an excel file to measure the differences in the distribution between the control and treated samples.

To quantify viroplasm formation, cells exhibiting pericentriolar aggregates were visually scored. Cells were considered positive when viroplasm were observed in close proximity to the centrosomes, with centrosomal localization serving as the reference criterion. For estimation of viroplasm size, the Feret’s diameter was measured using the Fiji software. The radius of each viroplasm was then calculated by dividing the Feret’s diameter by two. To analyse the centrosome amplification the centrosomes were marked with the pericentrosomal matrix marker γ-tubulin. We quantified all the three populations including the cells with acentriolar population (0), cells with 1 or 2 centrosomes (normal) and >2 centrosomes (supernumerary) indicating centrosome amplification. For the MTOC functional scoring the microtubule re-nucleation was scored by counting the number of asters appearing in the total population across mock in comparison to the JEV infected cells using α-tubulin staining pattern in the confocal micrographs acquired 1 min post release from cold shock. Similarly, the microtubule cytoskeleton formation status was assessed by the appearance of cytoplasmic tubulin fibres in the 4 min post release time point and the same were counted manually for quantification.

For densitometric analysis of immunoblots, the signal intensity of the protein of interest was quantified in Fiji using a constant region of interest (ROI) across all lanes with a fixed area for each lane. Background intensity was measured separately for the same size of area and subtracted from the raw signal to obtain normalized protein intensity values. The intensity of the corresponding loading control was then quantified and used for normalization. Finally, the expression level of the protein of interest was calculated as fold change relative to the appropriate experimental reference condition.

To estimate the polarity of VPs, the nucleus, centrosomes, and VPs were defined by setting the background threshold, and the centroid of the nucleus and the centre of mass for the centrosome and VPs were identified. The reference axis was set using the nuclear centroid and the centre of mass of centrosome. Finally, the position of each VP was determined with respect to the reference axis using the angle formed by the centroid of the nucleus and the VPs centre of mass. These angles were then used to interpret the polarity of VPs with respect to the centrosomes, where 0° to 45°was considered aligned (strong), 45° to 90°was considered scattered with no alignment (none), and 90° to 180° was considered opposing polarity (opposite).

### Statistical Analysis

All datasets were collected from at least three independent biological experiments tabulated in excel file were imported and analysed statistically using GraphPad Prism 10 software. The P values were calculated using unpaired Student’s *t* test for two given grouped events, while one-way ANOVA test was performed for differences across more than two sample groups. Data distribution was considered to be normal for parametric Student’s *t* test and ANOVA test. One-way ANOVA Dunnet’s multiple comparison test was performed for all the plaque assay end point titer analysis conducted in this study. The significance is indicated by * for P≤0.01; ** for P≤0.001; *** for P≤0.0001 and **** for P≤0.00001.

## Online Supplemental Material

Figure S1 (related to Figure 1) shows the validation of centrosome localization in HeLa cells (Figure S1A) and the expression of respective proteins through western blot analysis (Figure S1B-E).

Figure S2 (related to Figure 2) shows the expression levels of JEV NS3-HA with and without proteasomal inhibition (Figure S1A-B). It also compares the field views showing pattern of distribution of NS3 between JEV infection and JEV NS3-HA overexpression (Figure S1C).

Figure S3 (related to Figure 3) exhibits the 3D-SIM video and snapshots of JEV viroplasm around centrosomes enclosed by vimentin cage (Figure S3A-B). Confocal micrographs representing the number of viroplasms for the indicated time points (Figure S3C, related to Figure 3H).

Figure S4 (related to Figure 6) shows the effect of 24 hr Centrinone treatment in the infected population with no difference between DMSO and Centrinone treatment at 24 hpi.

Table S1 and S2 provides information about the cloning and mutagenic primers respectively. Table S3 lists the antibodies used in this study.

## Data availability statement

All data underlying the research presented in the manuscript are included either in the article or the supplementary data as cited in the manuscript text.

## Acknowledgement

We would like to acknowledge Prof. Sudhanshu Vrati, for gifting the Japanese Encephalitis Virus (strain <u>P20778</u>, GenBank accession no. <u>AF080251</u>) and Dr. Manjula Kalia for sharing the JEV NS3-HA expressing clone. pLV_Zika_NS3_Flag was a gift from Vaithi Arumugaswami (Addgene plasmid # 79635; http://n2t.net/addgene: 79635; RRID: Addgene_79635). bNS2B47NS3 was a gift from Dahai Luo (Addgene plasmid # 155320; http://n2t.net/addgene:155320; RRID: Addgene_155320). pCMV-Tag1-NS3 was a gift from Xin Wang (Addgene plasmid # 17645; http://n2t.net/addgene:17645; RRID:Addgene_17645). pSNAPf-C1 was a gift from Michael Davidson (Addgene plasmid # 58186; http://n2t.net/addgene:58186; RRID:Addgene_58186). We also thank Dr. Sudhanshu Vrati and Dr. Manjula Kalia for their critical reading and constructive comments on our manuscript.

## Funding

K.D. acknowledges RCB, Faridabad core funding aided by the Department of Biotechnology (DBT), Ministry of Science and Technology, India, for this work. K.D. is partly also supported by Ramalingaswami Re-entry Fellowship (BT/RLF/Re-entry/47/2020) from the Department of Biotechnology (DBT), Ministry of Science and Technology, India and partly by the Start-up Research Grant (SRG/2021/001466) from the Science and Engineering Research Board (SERB), Department of Science and Technology (DST), India. H.A. and M.Y. thank UGC, Government of India, for their student fellowship support during this study.

## Conflict of interest

The author(s) declared no potential conflicts of interest with respect to the research, authorship, and/or publication of this article.

## Generative AI statement

The authors declare that no Generative AI was used in the creation of this manuscript.

## Authors Contributions

K.D. Conceptualised, conceived, planned and performed experiments, analysed, interpret, written and edited the manuscript. H.A. planned and performed experiments, quantified and analysed the data, prepared the figures and edited the manuscript. P.D. has performed the high resolution imaging based assays, and the microtubule nucleation assay while M.Y. performed the plaque assays reported in this study while B.B. cultured and prepared the JEV virus used in this study. S.V. helped in the interpreting the antiviral studies and editing the manuscript.

## Supplementary figures

**Figure S1:**
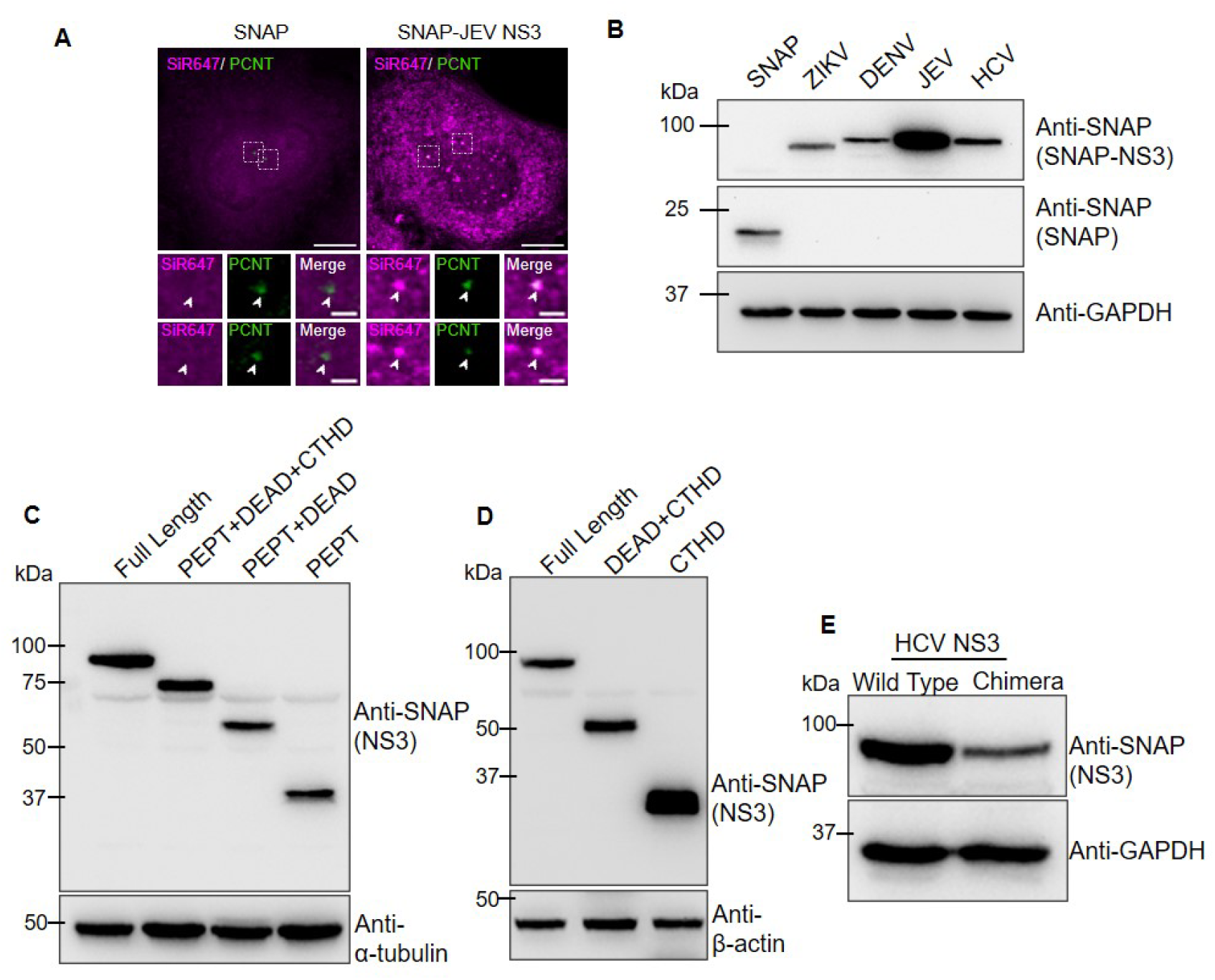
**(A)** Confocal micrograph representing the expression of SNAP only and SNAP-JEV NS3 (magenta) in HeLa-M cells. Centrosomes marked by Pericentrin (green) (Scale bar-10 μm). Zoom in Inset with arrow heads pointing centrosome (Scale bar- 2 μm). **(B)** Immunoblots showing the expression of SNAP tagged Flaviviral helicases (SNAP-NS3) along with the empty vector expressing SNAP alone (SNAP) in HEK293T cells using anti SNAP antibodies. Loading control-GAPDH. **(C)** Immunoblots showing the expression of C-terminal truncated JEV helicase constructs in HeLa-M cells. Loading control-α-Tubulin. **(D)** Western blot showing the expression of N-terminal truncated JEV helicase constructs in HeLa-M cells. Loading control-β-Actin. **(E)** Immunoblots showing the expression of wild type HCV NS3 along with the chimeric HCV construct where CTHD domain of HCV was replaced with the JEV NS3-CTHD domain. Loading control-GAPDH.

**Figure S2:**
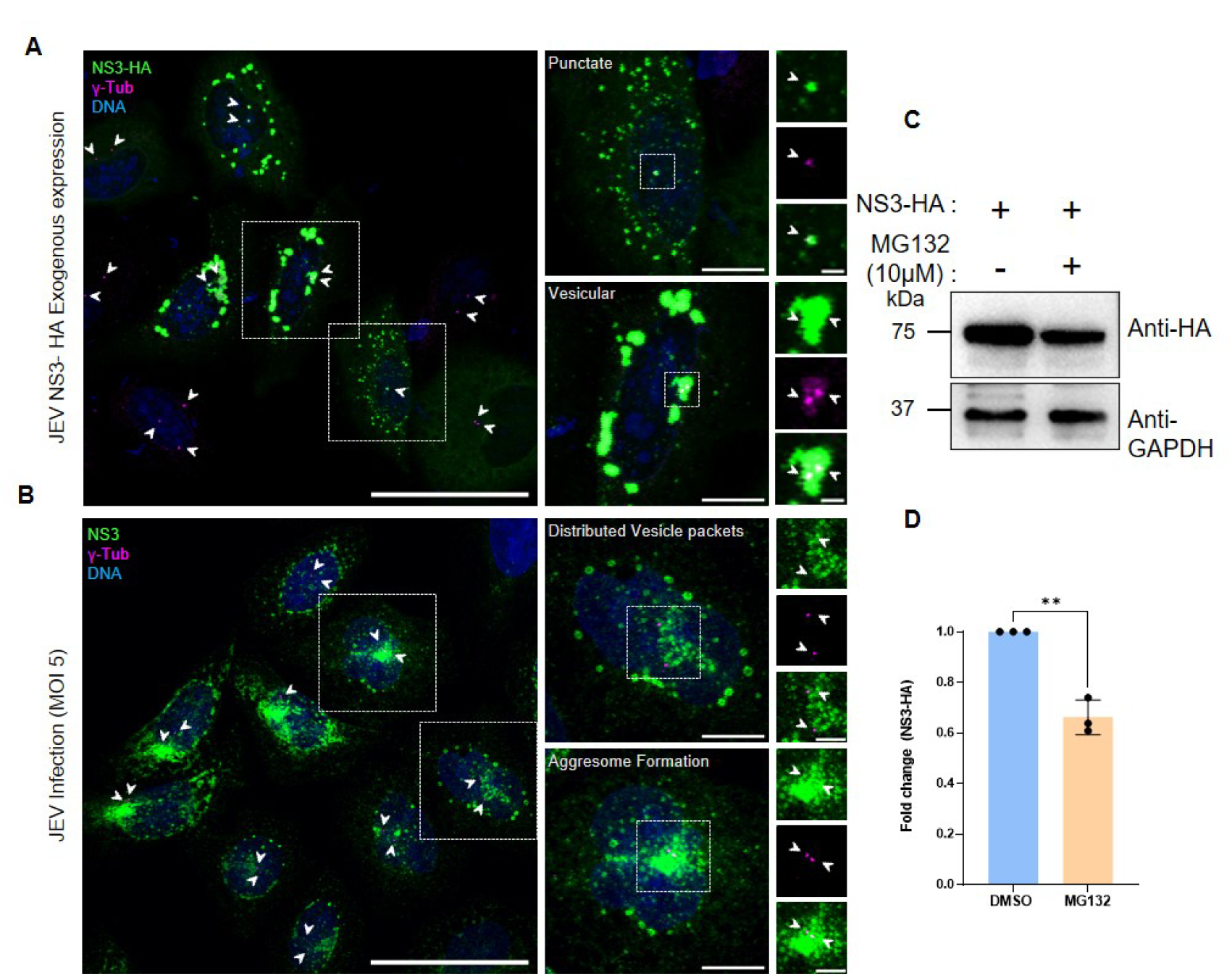
**(A)** HeLa-M cells exhibiting different localization patterns in exogenously expressed JEV-NS3 with HA tag (NS3-HA) visualized using anti-HA antibody are represented in the confocal micrographs (Scale bar: Field view -50 μm, single cell -10 μm, Zoom in inset -2 μm). **(B)** Similar distribution pattern of the anti-NS3 stained structures in JEV infected HeLa-M cells shows either distributed vesicle packets or aggresomes like perinuclear structures (Field view Scale bar- 50μm, Single cell Scale bar- 10μm, Zoomed in inset -5μm). NS3 (green), γ-tubulin (magenta) and DNA (cyan). White arrowheads indicate centrosome. **(C)** Immunoblots showing the expression of NS3-HA in DMSO and MG132 treated HeLa-M cells. GAPDH was used as the loading control. **(D)** Densitometric quantification of NS3-HA expression in DMSO and MG132 treated HeLa-M cell lysates (p value- 0.0011, **, N=3).

**Figure S3:**
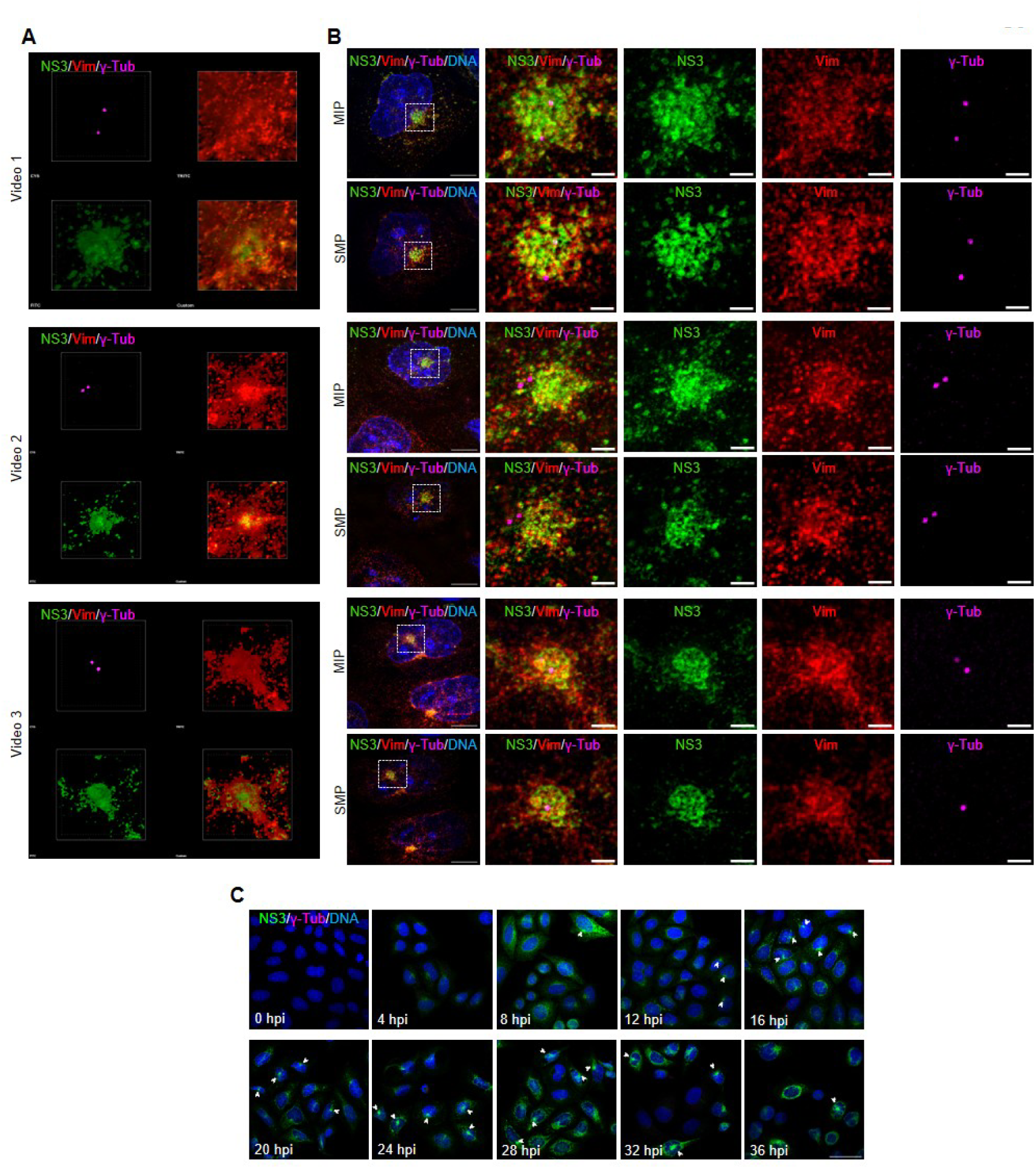
**(A)** Video files of three individual pericentriolar JEV viroplasm stained with anti-NS3 in green, anti-γ-tubulin in magenta and anti-vimentin in red. **(B)** 3D-SIM micrograph representing the maximum intensity projection (MIP) view and the single mid plane (SMP) view of the corresponding viroplasm appearing in panel A stained with anti-NS3 in green and anti-γ-tubulin in magenta, anti-vimentin in red and DNA in cyan stained with Hoechst. (Scale bar- 10 μm, Zoom in inset- 2 μm). **(C)** Confocal micrographs of HeLa-M cells infected with 5-MOI JEV for the indicated hpi tracing the viroplasm using anti-JEV NS3 antibody (green) along with the centrosomal marker γ-tubulin (magenta) and DNA using Hoechst (Scale bar- 10 μm). White arrowheads indicate pericentriolar viroplasm.

**Figure S4:**
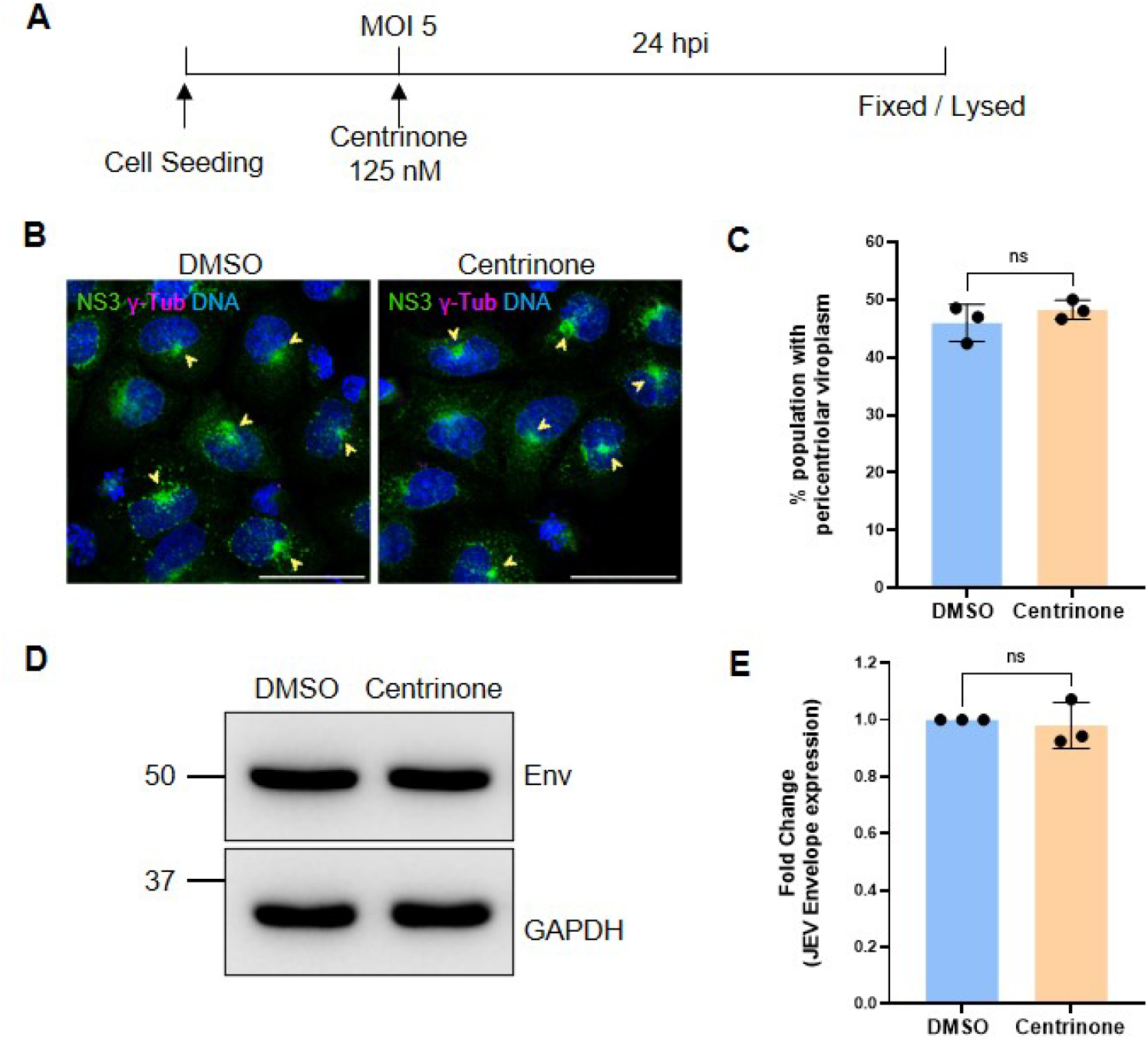
**(A)** Schematic of JEV infection with 125 nM Centrinone treatment for 24 hrs immediately after 5-MOI JEV infection. **(B)** Confocal micrographs showing JEV viroplasm (yellow arrowheads) in DMSO and Centrinone treated cells. NS3 is marked in green and γ-tubulin in magenta (Scale bar- 50 μm). **(C)** Quantification of percent infected population with pericentriolar viroplasm across DMSO vs Centrinone treated cells (p value- 0.3378, ns, N=3, n≥100). **(D)** Western blot analysis of DMSO vs Centrinone treated lysate using JEV Envelope antibody with GAPDH as loading control. **(E)** The same is quantified using densitometric analysis (p value- 0.6874, ns, N=3)

**Supplementary Table S1:**
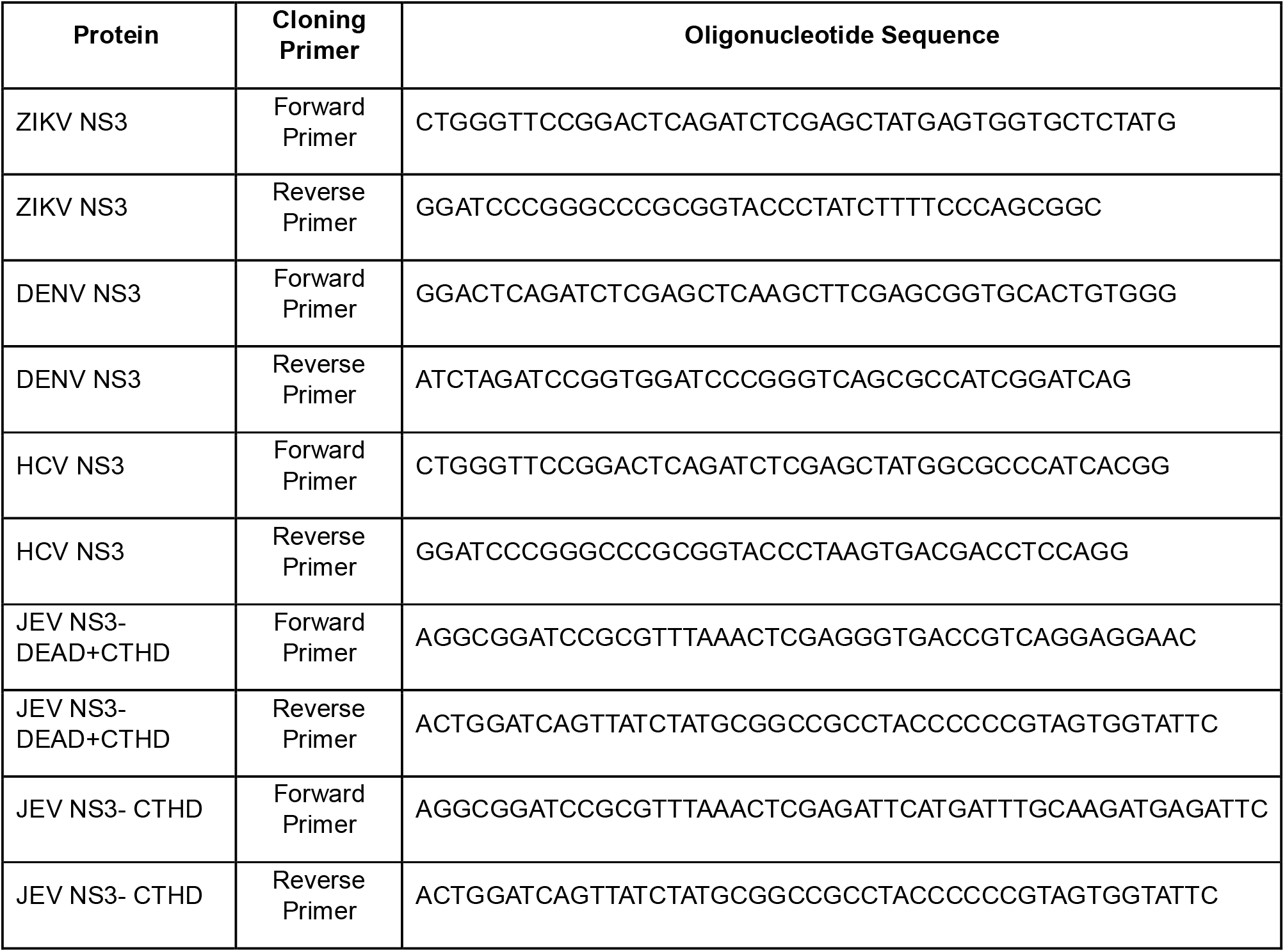
List of cloning primers.

**Supplementary Table S2:**
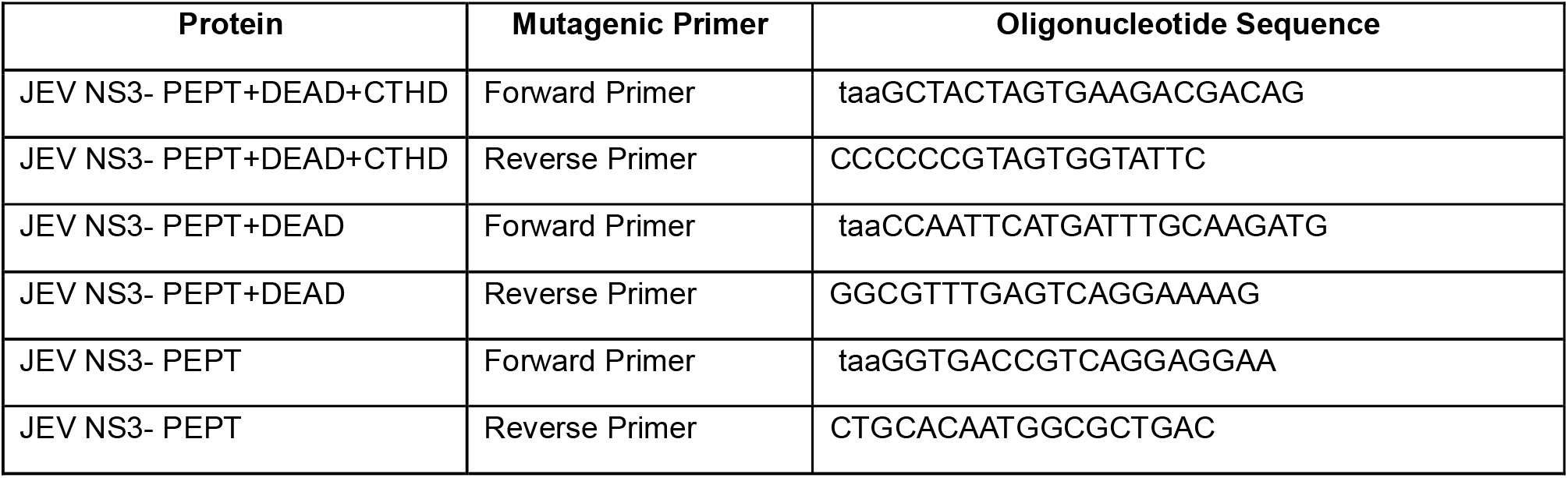
List of Mutagenic primers.

**Supplementary Table S3:** List of antibodies used for immunofluorescence (IF) staining and western blotting (WB) analysis.

| Antibody | Source | Cat. Number | Host Species | Dilution |
| --- | --- | --- | --- | --- |
| Anti-Pericentrin | Abcam | ab4448 | Rabbit | 1:4000 (IF) |
| Anti-γ-tubulin (GTU 88) | Sigma-Aldrich | T5326 | Mouse | 1:1000 (IF) |
| Anti-PCM1 | Sigma-Aldrich | HPA023370 | Rabbit | 1:500 (IF) |
| Anti-CEP152 | Invitrogen | MA518286 | Mouse | 1:200 (IF) |
| Anti-α-tubulin, AF-488 conjugate | Invitrogen | 322588 | Mouse | 1:500 (IF) |
| Anti-Vimentin (D21H3), AF-555 conjugate | Cell Signalling Technology | 9855S | Rabbit | 1:50 (IF) |
| Anti-Hemagglutinin (HA) | Bio Bharati Life Science | BB-AB0050A | Rabbit | 1:500 (IF) 1:1000 (WB) |
| Anti-JEV NS3 | GeneTex | GTX638821 | Rabbit | 1:800 (IF) |
| Anti-JEV NS5 | GeneTex | GTX131359 | Rabbit | 1:500 (IF) |
| Anti-JEV Capsid | GeneTex | GTX131368 | Rabbit | 1:100 (IF) |
| Anti-JEV Envelope | GeneTex | GTX125867 | Rabbit | 1:100 (IF) 1:5000 (WB) |
| Anti-dsRNA (J2) | Cell Signalling Technology | 76651L | Mouse | 1:800 (IF) |
| Anti-SNAP | Invitrogen | CAB4255 | Rabbit | 1:3000 (WB) |
| Anti-α-tubulin | Cell Signalling Technology | 3873S | Mouse | 1:10000 (WB) |
| Anti-GAPDH | Bio Bharati Life Science | BB-AB0060 | Rabbit | 1:2000 (WB) |
| Anti-β-actin | Cell Signalling Technology | 3700S | Mouse | 1:10000 (WB) |
| AF488 goat anti Mouse IgG (H+L) | Invitrogen | A11029 | Goat | 1:2000 (IF) |
| AF488 goat anti Rabbit IgG (H+L) | Invitrogen | A11034 | Goat | 1:2000 (IF) |
| AF568 goat anti Mouse IgG (H+L) | Invitrogen | A11004 | Goat | 1:2000 (IF) |
| AF568 goat anti Rabbit IgG (H+L) | Invitrogen | A11011 | Goat | 1:2000 (IF) |
| AF647 goat anti Mouse IgG (H+L) | Invitrogen | A21235 | Goat | 1:2000 (IF) |
| AF647 goat anti Rabbit IgG (H+L) | Invitrogen | A21245 | Goat | 1:2000 (IF) |
| Goat anti Rabbit IgG, HRP conjugate | Bio Bharati Life Science | BB-SAB 01A | Goat | 1:3000 (WB) |
| Goat anti Mouse IgG, HRP conjugate | Bio Bharati Life Science | BB-SAB 02A | Goat | 1:3000 (WB) |

## Notes

### Competing Interest Statement

The authors have declared no competing interest.

### Summary of Updates

A new result (Figure 5) comparing the viroplasm organization and centrosomal phenotypes between the non-neuronal and neuronal cell subtype is added. The MTOC status of neuronal and non-neuronal cells is responsible for the particular centrosomal phenotypes in these instances. The middle name of author Pranay Dudhe is now included in the list of authors, appearing as Pranay Eknath Dudhe. Supplemental Figure S4 is now modified. Figure S4A and S4B showing the integrated density of CEP152 post JEV infection is now added to the main figure panel as Figure 4C and 4D. A separate section on the limitations of the study has now been incorporated.

